# Deciphering neuronal variability across states reveals dynamic sensory encoding

**DOI:** 10.1101/2024.04.03.587408

**Authors:** Shailaja Akella, Peter Ledochowitsch, Joshua H. Siegle, Hannah Belski, Daniel Denman, Michael A. Buice, Severine Durand, Christof Koch, Shawn R. Olsen, Xiaoxuan Jia

## Abstract

Influenced by factors such as brain states and behavior, neurons exhibit substantial response variability even to identical stimuli. Because these factors are non-stationary, they dynamically impact the fidelity of sensory processing. However, it remains unclear how their relative impact on neuronal variability evolves over time. To address this question, we designed an encoding model with latent states to partition visual cortical variability across three crucial categories of sources: internal brain dynamics, behavior, and external visual stimulus. Applying a hidden Markov model to the rhythmic patterns of cortical local field potentials, we consistently identified three distinct oscillation states. Each state revealed a unique variability profile and a consistent descending trend of stimulus modulation across the visual hierarchy. Regression models within each state revealed a dynamic composition of factors contributing to the observed spiking variability, with the primary influencing factor switching within seconds. In the state dominated by high-frequency oscillations, sensory inputs and behavior exerted the most influence on population dynamics. Conversely, internal brain activity explained most of the variance in the state dominated by low-frequency oscillations. This heterogeneity across states underscores the importance of partitioning variability over time, particularly when considering the dynamic influence of non-stationary factors on sensory processing.

## 1 Introduction

The amount of information a sensory neuron carries about external stimuli is reflected in its repeated activity pattern in response to the same stimuli (Reinagel and Reid 2000). However, trial-to-trial variability, ubiquitous in the nervous systems (Shadlen and Newsome 1998), constrains the amount of sensory information in single-trial neural responses to the stimulus. It follows that the time course of this variance mimics the highly non-stationary dynamics of the underlying neuronal processes (Churchland et al. 2011, Churchland et al. 2010). For example, when animals actively explore their environment, the sensory cortex shows desynchronized responses in a manner that increases their responsiveness to stimuli (Poulet and Petersen 2008). Conversely, during periods of sleep or quiet wakefulness, cortical neurons tend to synchronize their activity, resulting in decreased sensitivity to external stimuli (White et al. 2012). Dissecting these non-stationary dynamics is critical to comprehending their role in information encoding and ultimately, perception.

Even with well-controlled experiments and behavior-monitoring techniques (Nath et al. 2019; Pereira et al. 2022), understanding how neuronal variability changes over time is challenging (Festa et al. 2021). This is further complicated by the high-dimensional interactions between the various sources of neuronal variability: external stimuli, behavior, and internal brain dynamics (Goris et al. 2014). To address this complexity, a common strategy involves the identification of meaningful temporal patterns and potential latent variables that can capture the evolving dynamics of neural activity. These patterns, which accurately capture the internal brain dynamics, are typically referred to as “brain states” (Harris and Thiele 2011; McGinley et al. 2015; Poulet and Petersen 2008; Recanatesi et al. 2022).

Brain states, characterized by distinct patterns of neural activity and functional connectivity, play a pivotal role in shaping the dynamics of neuronal variability (Recanatesi et al. 2022; White et al. 2012), influencing how sensory information is processed (Churchland et al. 2010; Lombardo et al. 2018) and behaviors are executed (McGinley et al. 2015; Poulet and Petersen 2008). For instance, during heightened attention, decreases in the correlations between the trial-to-trial fluctuations in the responses of pairs of neurons, serve to enhance the signal-to-noise ratio of the entire population, improving behaviors (Cohen and Maunsell 2009). Likewise, several studies have shown that random fluctuations in the processing of sensory stimuli originate from rapid shifts in the animal’s arousal state (Britten et al. 1996; McGinley et al. 2015). Tightly linking internal brain dynamics to behavior, brain states serve as an ideal temporal framework to study the non-stationarity of neuronal variability.

Recently, researchers have leveraged advanced machine-learning tools to explain single-trial neural activity by incorporating extensive stimulus and behavioral features (Musall et al. 2019; Pandarinath et al. 2018; Stringer et al. 2019). While these studies reveal the multi-dimensional nature of neuronal variability, they often assume that neuronal variability remains constant over time. To address this gap, several parallel lines of research have used latent dynamical models to study the temporal patterns of neuronal variability (Ashwood et al. 2022; Calhoun et al. 2019; Poulet and Petersen 2008; Recanatesi et al. 2022). However, these studies have not explicitly explored the different sources contributing to variability, as it changes over time. Consequently, our understanding of how various sources dynamically contribute to the non-stationarity of neuronal variability remains limited (Figure 1**A**).

Here, we present a comprehensive investigation of how internal and external factors collectively shape the time course of neuronal variability to influence sensory coding. We used the Allen Brain Observatory Visual Coding dataset, which comprises simultaneous recordings of local field potentials (LFPs) and spiking activity from hundreds of Neuropixels channels in multiple visual areas along the anatomical hierarchy (Harris and Thiele 2011). As mice passively viewed natural movies, we applied Hidden Markov Models (HMMs) (Rabiner 1989) on LFP data extracted from six visual cortical regions to establish a global temporal framework of internal latent states. Quantifying various aspects of variability across individual trials and neuronal populations, we uncovered significant non-stationarity in neuronal variability across states. These findings indicated dynamic changes in the efficiency of sensory processing over time, revealing a consistent descending trend of stimulus induced variability across the visual hierarchy. To elucidate the relationship between these non-stationarities and various sources of variability, we designed a novel HMM-based encoding framework to partition variability across three crucial factors: internal brain dynamics, spontaneous behavior, and external visual stimuli. Through this model, we quantified the time-varying contributions of these sources to single-trial neuronal and population dynamics. We found that even during persistent sensory drive, neurons dramatically changed the degree to which they were impacted by sensory and non-sensory factors within seconds. Taken together, our results provide compelling evidence for the dynamic nature of sensory processing, while emphasizing the role of latent internal states as a dynamic backbone of neural coding.

## 2 Results

We analyzed the publicly available Allen Brain Observatory Neuropixels dataset, previously released by the Allen Institute (Siegle, Jia, et al., 2021). This dataset comprises simultaneous recordings of spiking activity and local field potentials (LFPs) from six interconnected areas in the visual cortex of mice (n = 25) passively viewing a variety of natural and artificial visual stimuli (Figure 1**B**). To estimate the dynamic nature of internal state fluctuation during sensory processing, we focused our analysis on data recorded during repeated presentations of a 30-second natural movie. We used a continuous stimulus to mitigate sudden transients in activity induced by abrupt changes in the visual stimuli. Lastly, the application of quality control metrics yielded, on average, 304 *±* 83 (mean *±* std) simultaneously recorded neurons distributed across layers and areas per mouse (see Methods).

Previous studies (Siegle, Jia, et al., 2021, Jia et al. 2022) demonstrated that the functional hierarchy of visual areas aligns with their anatomical organization (Harris et al. 2019). This hierarchy places the primary visual cortex (V1) at the bottom, followed by rostrolateral (RL), lateromedial (LM), anterolateral (AL), posteromedial (PM), and anteromedial (AM) areas (Figure 1**C**). Here, we consider this visual hierarchy as a first-order approximation of signal processing stages to study signal propagation and information encoding while crucially accounting for the non-stationarity in spiking variability that arises due to influences from fluctuating internal and external factors.

### Identification of oscillation states from local field potentials

Internal brain states can vary without clear external markers, making their quantification challenging. To capture state changes associated with internal processes, we employ a definition of brain states derived using LFPs recorded invasively from six visual areas (Siegle, Jia, et al., 2021). LFPs reflect aggregated sub-threshold neural activity and capture the highly dynamic flow of information across brain networks (Buzśaki et al. 2012). The spectral decomposition of LFPs reveals different frequency bands that correlate with specific cognitive states (Berens et al. 2010; Caton 1875; Jacinto et al. 2013), sensory processing (Akella et al. 2021; Di et al. 1990; Jia and Kohn 2011; Schroeder et al. 2001; Victor et al. 1994), and behavior (DeCoteau et al. 2007; Murthy and Fetz 1996; Scherberger et al. 2005). We found that LFPs in the mouse visual areas also revealed a distinct frequency spectrum across time, whose dynamics were strongly coupled to arousal-related behavioral variables (Figure 2**A**). Accordingly, we envisioned that a latent state model could reflect the underlying latent brain dynamics by capturing the dynamic patterns of the LFP spectrum, such that each latent state reflects an oscillation state. To extract these oscillation states from LFPs in the visual area, we employ Hidden Markov modeling (Beron et al. 2022; Linderman et al. 2017; Rabiner 1989) on filtered envelopes of LFPs within distinct frequency bands: 3 - 8 Hz (theta), 10 - 30 Hz (beta), 30 - 50 Hz (low gamma), and 50 - 80 Hz (high gamma). This approach enabled us to fully capture LFP power across the 3 *−* 80 Hz frequency range (Figure S1**A**), while also aligning with the observed frequency boundaries in the spectral decomposition of LFPs (Figure 2**B**, left panel). Finally, to capture laminar dependencies, the overall input to the HMM also comprised LFPs from superficial, middle and deep layers in all visual areas (one channel each from layer 2/3, layer 4, layer 5/6; Figures 2**B** (middle panel), S1**E, F**).

**Figure 1:**
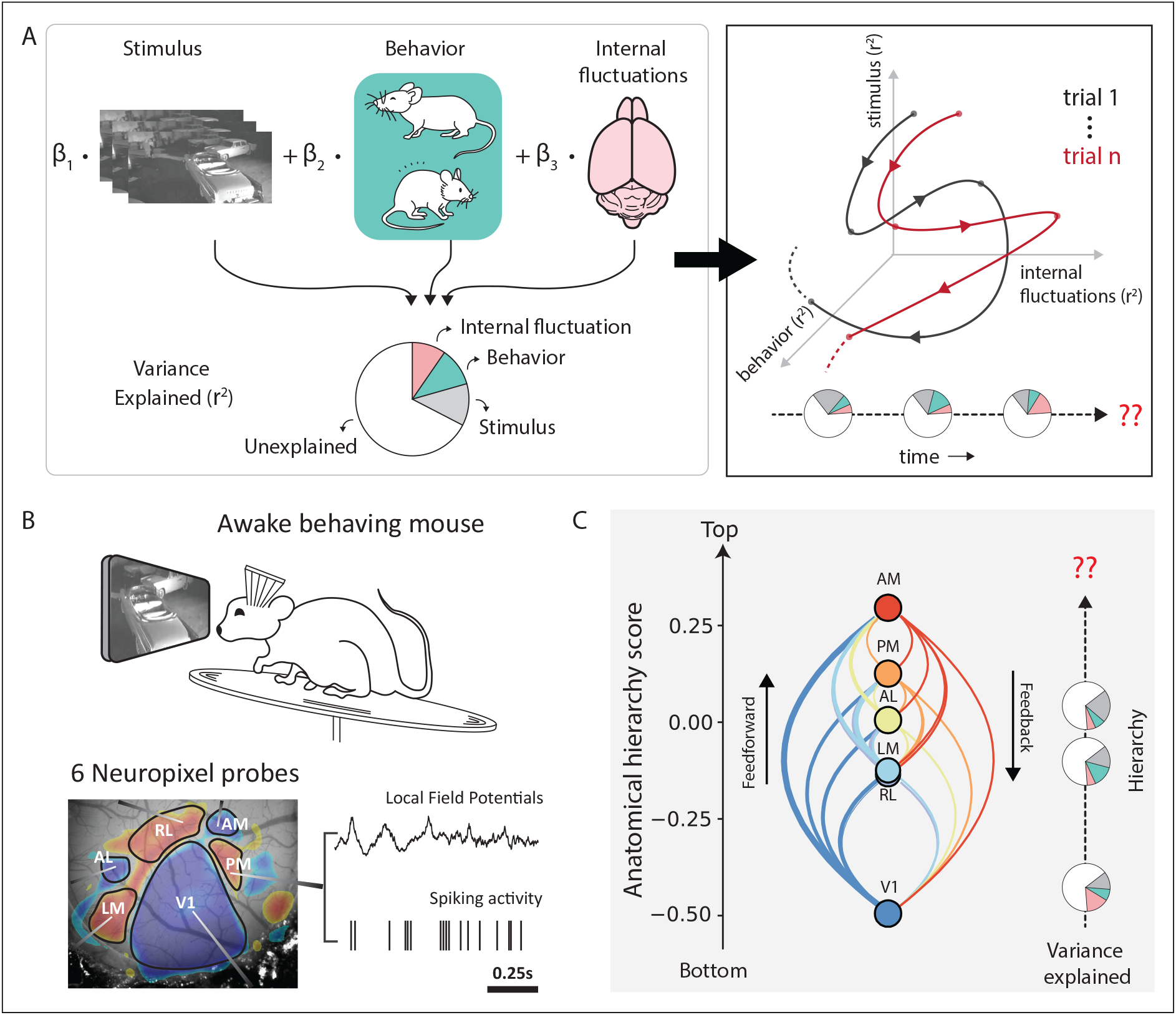
Schematic overview on deciphering variability across time and hierarchy. **A**, Neuronal variability is a combined effect of influences from independent stochastic processes including external sensory factors, behavior, and fluctuations in internal brain states. The resulting neuronal responses exhibit a variable temporal structure across trials and individual neurons. Capturing these temporal dynamics is a challenging problem and lies at the core of understanding the functional role of neuronal variability. **B**, Top: Schematic of the experimental setup. Bottom: Neuropixels probes in six visual cortical areas simultaneously record local field potentials and spiking activity. A retinotopic sign map overlaid on the vasculature image guides area-specific targeting. **C**, Anatomical hierarchy scores of the six visual areas recomputed from (Harris et al. 2019). Studying variability along the visual hierarchy can reveal important insights about information propagation and encoding at each stage of signal processing.

We found that LFP dynamics in the visual cortex consistently unfolded through three reliable oscillation states across all mice (see Methods; Figure 2**C**, 3.08 *±* 0.39, n = 25 mice, mean *±* std). These states did not depend on stimulus types (Figure S4**A, B**), specific visual areas (Figure S1**B, C**), or layers (Figure S1**E, F**). The identity of the inferred states was also remarkably consistent across mice, each characterized by a distinct distribution of the power spectrum: a high-frequency state (*S_H_*), a low-frequency state (*S_L_*), and an intermediate state (*S_I_*). While the high-frequency state is characterized by increased power in the low and high gamma bands, slow oscillations dominate the low-frequency state dynamics in the theta frequency ranges (Figures 2**D****, E**, S2**C**). LFP power distribution in the intermediate state is more uniform.

**Figure 2:**
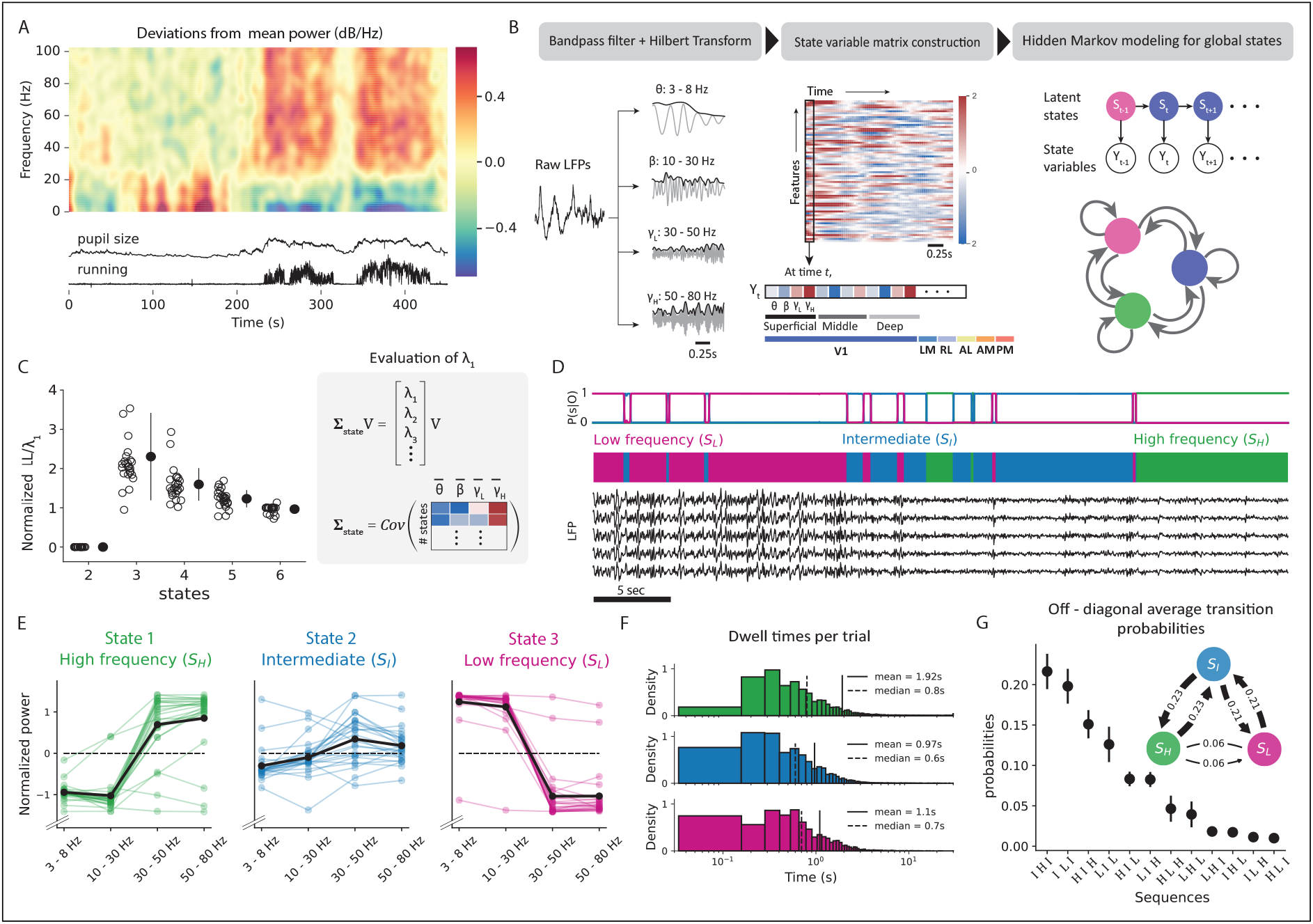
Properties of internal oscillation states identified from local field potentials in awake behaving mice. **A**, Top: LFP power modulations in V1 recorded from mice passively viewing a naturalistic movie. Bottom: Time course of running speed and pupil area during the same time period. **B**, Schematic to identify oscillation states using local field potentials. Discrete states are defined based on frequency-specific transients of LFPs from 6 visual areas. Hidden Markov model (HMM) uses Hilbert transforms in the theta (3-8 Hz), beta (10-30 Hz), lower gamma (30-50 Hz), and higher gamma (50-80 Hz) frequency ranges. **C**, Left: Model comparison among HMMs over a range of latent states using three-fold cross-validation. Test set log-likelihood penalized by state similarity (*λ*_1_) is reported. Right: Evaluation of state similarity (*λ*_1_) as the top eigenvalue of the HMM covariance matrix. **D**, Top: State posterior probabilities identified by the HMM. Bottom: LFPs from V1 alongside their respective latent states in the same duration. **E**, LFP power distribution in the three-state model. In state-1, or the high-frequency state, LFPs are dominated by high-frequency gamma oscillations. State 3, or the low-frequency state, has characteristic slow oscillations in the theta band. **F**, Histogram of state dwell times in each trial across all states and all mice. **G**, Average probability of observing 3-step or 2-step (inset) transition sequences to different states. Transition probabilities were calculated from observed sequences averaged across all mice.

These oscillation states demonstrate stable dynamics, as reflected by the large values along the diagonal of the transition matrix, ranging between 0.94 and 0.99 (Figure S3**B**). Dwell time in a state averaged around 1.5 *±* 0.14s (mean *±* sem, n = 3 states) (Figure 2**D****, F**), and the transition intervals between consecutive states (the interval around a transition during which the HMM posterior probability is *<* 80 %) were significantly shorter than the dwell times, lasting only for about 0.13 *±* 0.006s (mean *±* sem). Additionally, direct transitions between the low- and high-frequency states were rare and required transitioning through the intermediate state, as evident in both two- and three-step transition sequence-probability trends (Figure 2**G**). Consequently, mice spent only short durations in the intermediate state (0.97 *±* 0.001s, mean *±* sem), while they spent the most prolonged durations in the high-frequency state (1.92 *±* 0.003s, mean *±* sem, 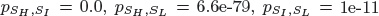, n = 25 mice). Notably, this state property was dependent on stimulus type (Figure S4**D**). During repeated presentations of the drifting grating stimulus, transitions between the extreme states of low- and high-frequency were much faster and more likely (Figure S4**E,F**). This significantly reduced the amount of time mice spent in the intermediate state (0.25 *±* 0.0001s, p = 0, Figure S4**C**). However, in the absence of any stimulus, mice tended to spend longer durations in the intermediate state (1.16 *±* 0.001s, p = 3.5e-29). We attribute these differences to the strong neural responses evoked by sudden transitions of the visual stimulus such as, the onset and offset of drifting gratings stimuli.

**Figure 3:**
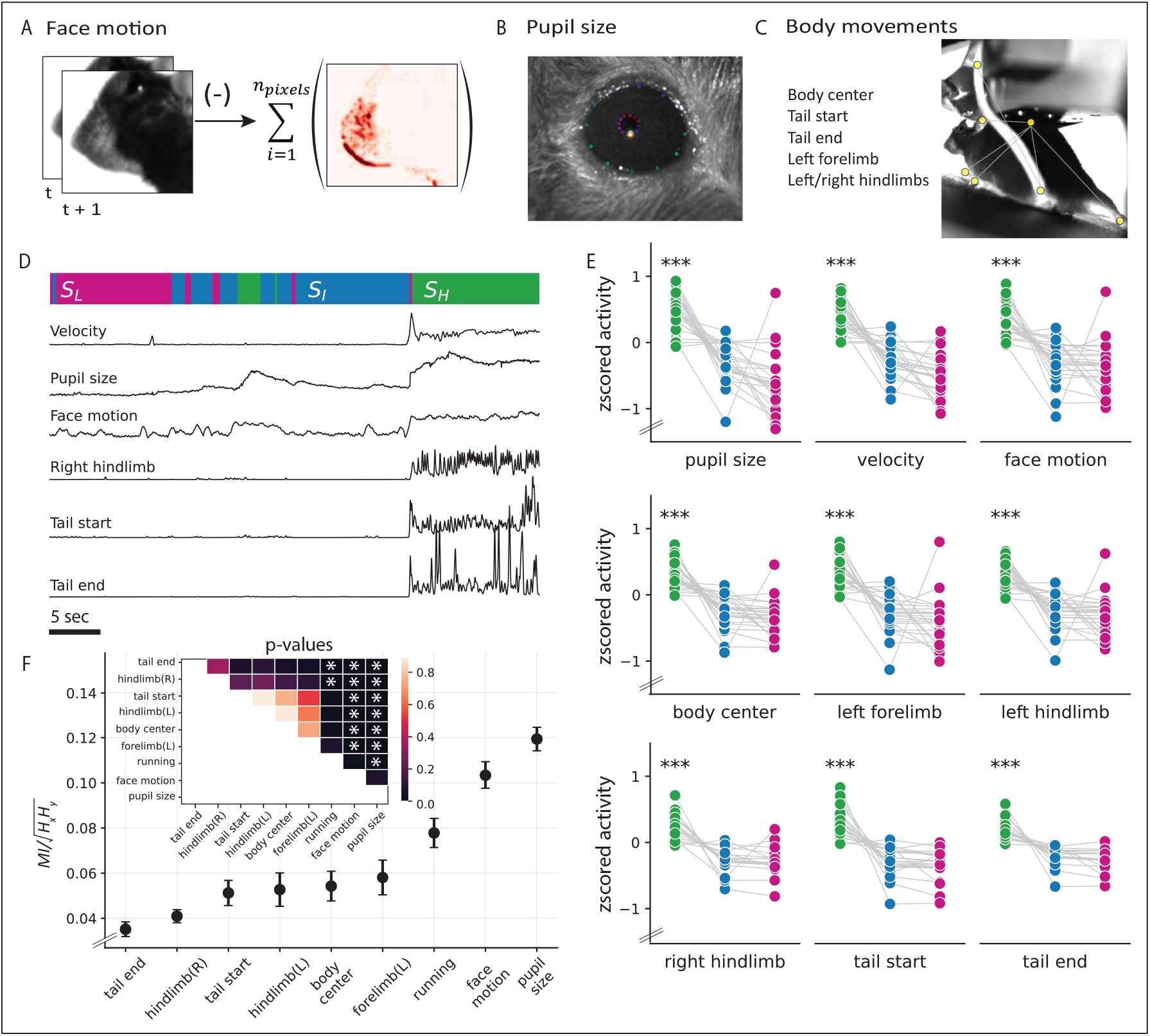
Behavioral correlates of the observed oscillation states. **A**, Face motion energy evaluated as the absolute value of the difference between consecutive frames. **B**, Eye and pupil tracking. Tracking points were identified using a universal tracking model trained in DeepLabCut. **C**, Animal pose estimation. Specific, visible body parts were tracked using a universal tracking model trained in SLEAP. **D**, Example snippet of behavioral changes alongside the animal’s current oscillation state. *S_H_* : High-frequency state (green), *S_I_* : Intermediate state (blue), and *S_L_*: Low-frequency state (pink). **E**, Comparison of the average movement of specific body parts across states (*p_S_H_,S_I_,L_*, pupil size: p = 2.8e-15, velocity: p = 2.0e-17, face motion: p = 6.3e-13, body center: p = 2.6e-18, left forelimb: p = 1.2e-13, left hindlimb: p = 4.9e-14, right hindlimb: p = 3.0e-11, tail start:, p = 3.0e-16, tail end: p = 2.0e-11, n = 25 mice). **F**, Mutual information (MI) between behavioral variables and the inferred HMM states (mean *±* sem, n = 25 mice).

### Correlation between oscillation states and body movements

Brain state variations often exhibit strong correlations with the animal’s behavioral context (Mccormick et al. 2020; Zagha and McCormick 2014). Indeed, several studies have reported neural activity changes in the visual cortex associated with various behavioral features (Bennett et al. 2013; Musall et al. 2019; Niell and Stryker 2010). To this end, we examined the behavioral correlates of the oscillation state patterns, comparing pupil size, running speed, and facial, limb, and tail movements across different states (Figure 3**A-C**). Our investigation revealed a strong association between behavioral movements and internal oscillation states across subjects (Figure 3**E**). Notably, a shift to the high-frequency state corresponded closely with increased movements and pupil size (Figure 3**D**), suggesting increased arousal levels in this state. Conversely, mice tended to be at rest in the low-frequency state while only making small movements in the intermediate state (McGinley et al. 2015; Reimer et al. 2016; Vinck et al. 2015).

Several studies have considered locomotion as an indicator of brain state to examine variations in visual encoding (Saleem et al. 2013; Stringer et al. 2019). To quantify the relationship between internal oscillation states and different behavioral features, we calculated the mutual information (MI) between the states and each behavioral feature (Sanchez Giraldo et al. 2015). We found that changes in the oscillation states were more faithfully mimicked by pupil size or facial movements (Figure 3**D**), reporting significantly higher MI than all other behavioral responses (*MI_pupil_* = 0.12 *±* 0.006*, MI_face_* = 0.1 *±* 0.006, mean *±* sem, n = 25 mice), including running (*MIr_unning_* = 0.08 *±* 0.007, mean *±* sem, n = 25 mice, Figure 3**F**). This held true despite the strong positive correlations between all behavior variables (r = 0.4 *±* 0.03, mean *±*, sem, n = 25 mice), and especially between running, facial movement, and pupil size (r = 0.6 *±* 0.04, mean *±*, sem, n = 25 mice). Importantly, all behaviors associated with running (movements in the proximal end of the tail, left limbs, and body center) reported similar MI with the oscillation states. To further validate these results, we used HMMs to quantify behavioral states in individual mice, fitting individual models to pupil size, face motion, and running measures. Upon comparing these behavioral states with oscillation states, stronger correlations emerged with pupil size and face motion than with running speed (Figure S5**B**; p = 0.0007, n = 25 mice). We attribute these differences to the dissociation between pupil size and running speed, particularly in cases where pupil dilation occurs, even when the mouse remains stationary (Figure S5**A**). These results suggest that facial movements serve as a reliable representation of the underlying internal states reflected in voluntary behavior, almost as good as the involuntary changes in pupil size (Crombie et al. 2024).

### Neuronal variability changes across oscillation states and visual hierarchy

After defining the internal oscillation states and establishing their relation to behavior and arousal state, we wondered how spiking variability changes across these states. Across states, we observed distinct variations in population activity and synchronization levels (Figure 4**A-C**). Consistent with previous observations of attentional effect (Cohen and Maunsell 2009), increased spiking activity (av. % increase = 7.7 *±* 1.6, mean *±* sem, p = 6.3e-5, n = 25 mice) and decreased correlation (av. % decrease = 36.6 *±* 3.4, mean *±* sem, p = 1.3e-10, n = 25 mice) were typical of the high-frequency state. Moreover, the transition-state-like properties of the intermediate state were broadly consistent across various neuronal properties (Figure 4**B****, C**) and behavior (Figure 3**E**). Bolstered by these findings, we evaluated three types of variability in single neurons to capture complementary aspects of neuronal variability: percentage of shared variance within a population, spike timing variability, and variability in spike counts across trials.

Previous studies have shown that variability shared within a neuronal population can constrain information propagation between processing stages (Averbeck et al. 2006; Denman and Reid 2019; Kohn et al. 2016; Lin et al. 2015). This is because shared variance within a population may not average out (Azeredo da Silveira and Rieke 2021; Moreno-Bote et al. 2014), leading to a deterioration of the population’s coding capacity. To study how shared variability evolves across various internal states, we used factor analysis (FA) (Williamson et al. 2016) to partition the spike count variability into its shared and independent components (Figure 4**D**, top). Within a neuronal population, the shared component quantifies co-fluctuations in firing rates among individual neurons, while the independent component captures their Poisson-like variability. Percentage of shared variability was then evaluated as the ratio between each neuron’s shared and total variance. Consistent with previous findings that noted more synchronization within a population during low-arousal states (Mccormick et al. 2020; Zagha and McCormick 2014), the percentage of shared variability was highest during the low-frequency state (Figure 4**D**, bottom). In this state, fewer factors influenced the observed patterns of variation compared to the other states (number of FA components, *S_H_* = 21 *±* 1, *S_I_* = 19 *±* 1, *S_L_* = 16 *±* 1, p = 1.8e-06). Neurons within V1 reported a larger shared component than neurons within other areas (Figure S6**A**). The percentage of shared variance decreased along the visual hierarchy in the high-frequency state, (Pearson correlation r = -0.85 with anatomical hierarchy score, p = 0.043, while the trends were not significant in the intermediate and low-frequency states (*S_L_* : Pearson’s r = -0.76, p = 0.08, *SI* : r = -0.59, p = 0.22). Compared to higher visual areas, neurons in early visual areas are known to be more modulated by the temporal features of visual stimuli (Matteucci et al. 2019, Siegle, Jia, et al., 2021). Thus, we attribute the observed decreasing trends to rapid variations in luminance or moving edges in the natural movie, that likely induce stronger temporally coherent activity within a population in lower visual areas than in higher visual areas.

To study variability in spike timing, we measured the histograms of inter-spike intervals (ISI) and their associated coefficients of variation (Softky and Koch 1993). Coefficient of variation (CV) of each neuron was evaluated as the ratio between the standard deviation and mean of the ISI distributions. Therefore, the farther a neuron’s CV deviates from 0, the more irregular the neuron’s firing (Figure 4**E**, top left). Evaluating CV in a state-specific manner, we found that neurons during the high-frequency state had broader ISI distributions than during other states (Figure 4**E**, top right), and accordingly, fired more irregularly in this state (Figure 4**E**, bottom). along the visual hierarchy, spike timing variability decreased irrespective of the internal state (Figure 4E, bottom, *S_H_* : Pearson’s r with anatomical hierarchy score = -0.93, p = 0.006; *S_I_* : Pearson’s r = -0.97, p = 0.001, *S_L_* : Pearson’s r = -0.93, p = 0.007). Consistent with our expectation that V1 neurons more faithfully represent the features of the time-varying visual stimuli (Chaudhuri et al. 2015; Matteucci et al. 2019; Murray et al. 2014, Siegle, Jia, et al., 2021), we found that activity of V1 neurons was the most irregular.

**Figure 4:**
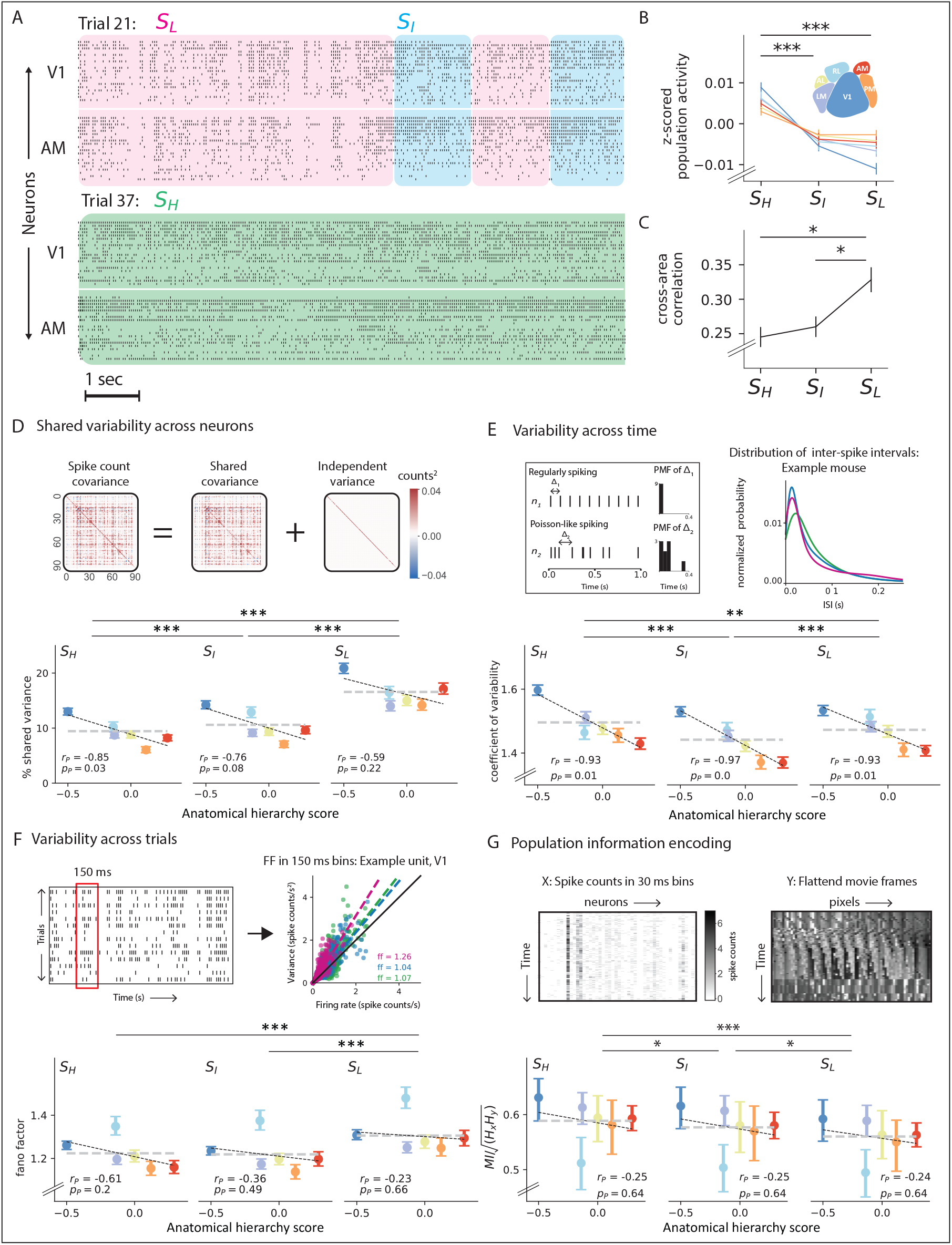
Neuronal variability and information encoding across states and the visual hierarchy. **A**, Raster plots (*∼* 10*s*) showing the response of 25 units, each from V1 and AM, during two trials in which the mouse was in different states. Each row represents the activity of the same single neuron across the two trials. *S_H_* : High-frequency state (green), *S_I_* : Intermediate state (blue), and *S_L_*: Low-frequency state (pink). **B**, State and area-specific population activity, z-scored and averaged across all mice (*pS_H_,S_I_* = 1.4e-05, *pS_H_,S_L_* = 3.0e-07, *pS_I_,S_L_* = 0.90, n = 25). **C**, Average pairwise correlation between averaged neuronal population activity in different visual areas as a function of oscillation states (*pS_H_,S_I_* = 1.5, *pS_H_,S_L_* = 0.002, *pS_I_,S_L_* = 0.002, n = 25). **D**, Population shared variance. Top: Separation of shared and independent variance using factor analysis (FA). FA partitions the spike count covariance matrix into shared and independent components. Bottom: Percent shared variance plotted against the anatomical hierarchy scores of the visual areas in each oscillation state, averaged across all units (*pS_H_,S_L_* = 0, *pS_I_,S_L_* = 5e-85, *pS_H_,S_I_* = 8.9e-6, n = 7609 units). **E**, Neuronal variability across time, quantified using the coefficient of variation (CV). Top-left: Simulated distributions of inter-spike-intervals (ISI) for regular and Poisson-like firing. For a very regular spike train, a narrow peak in the ISI histogram corresponds to CV *≈* 0, whereas Poisson-like variability in the spike trains leads to an exponentially distributed ISI histogram with CV = 1 Top-right: Distribution of ISIs in each oscillation state over a 2.5sec range. Bottom: CV along the visual hierarchy (quantified as anatomical hierarchy scores) and across oscillation states, averaged across all units (*pS_H_,S_I_* = 1.3e-17, *pS_H_,S_L_* = 7.1e-04, *pS_I_,S_L_* = 2.0e-06, n = 7609 units). **F**, Neuronal variability across trials, quantified using Fano factor (FF). Top-left: Evaluation of FF as an average of the FF ratio over non-overlapping windows of 150 ms with at least ten trials in each state Top-right: Mean spike count versus variance overall times in each state for an example cell in V1. Bottom: FF along the visual hierarchy and across brain states, averaged across all units (*pS_H_,S_L_* = 3.6e-15, *pS_I_,S_L_* = 1.8e-16, n = 7609 units). Pearson correlation with hierarchy scores excluding RL, *S_H_* : *r_p−RL_* = *−*0.93*, p_p−RL_* = 0.02; *S_I_* : *r_p−RL_* = *−*0.59*, p_p−RL_* = 0.3; *S_L_* : *r_p−RL_* = *−*0.35*, p_p−RL_* = 0.57 **G**, Information encoding along the visual hierarchy across all oscillation states, quantified using mutual information (MI). Top: For each trial, MI was evaluated between the population spike count matrix and a matrix of flattened movie frames at time points corresponding to each state using a matrix-based entropy estimator. Bottom: MI across the visual hierarchy and oscillation states averaged across all mice (*pS_H_,S_I_* = 0.03, *pS_H_,S_L_* = 2.2e-09, *pS_I_,S_L_* = 0.03, n = 25). Pearson correlation with hierarchy scores excluding RL, *S_H_* : *r_p−RL_* = *−*0.92*, p_p−RL_* = 0.03; *S_I_* : *r_p−RL_* = *−*0.89*, p_p−RL_* = 0.04; *S_L_* : *r_p−RL_* = *−*0.73*, p_p−RL_* = 0.15. Error bars in **D-G** represent 95% confidence intervals.

In visual system studies, trial-to-trial variability is commonly assessed using the Fano factor (FF, Fano 1947), which quantifies the ratio of variance to mean spike count across trials. An FF of 1 corresponds to a Poisson process, indicating that individual action potentials are generated randomly according to a constant firing rate. To ensure the relevance of our analysis to the visual stimulus, we evaluated FF of neurons with receptive field locations near the screen’s center (Kara et al. 2000; Softky and Koch 1993, see Methods, Figure 4**F**, top). Overall, single neurons in the visual cortex showed greater-than-Poisson variability with FF averaging around 1.47 *±* 0.6 (mean *±* std). Specifically, spike counts in the low-frequency state showed the largest trial-to-trial variability, suggesting it is less modulated by visual stimuli. In contrast, trial-wise variability was comparable across the intermediate and high-frequency states (Figure 4**F**, bottom). Interestingly, neurons in RL reported the highest variability across visual areas (Figure 4**F**, bottom), where regardless of the animal’s internal state, these neurons generally reported higher FF (Figure S6**C**). Accordingly, excluding area RL from the analysis revealed a decreasing trend in the trial-to-trial variability along the hierarchy in the high-frequency state (*S_H_* : Pearson’s r with anatomical hierarchy score = -0.93, p = 0.02; *SI* : Pearson’s r = -0.59, p = 0.3, *S_L_* : Pearson’s r = -0.35, p = 0.57).

Based on these results, we hypothesized that lower shared variance and trial-to-trial variability in spiking activity during the high-frequency state would improve stimulus encoding (Figure 4**D****, F**). Meanwhile, the increased spike timing variability during this state could be due to better encoding of the temporal changes in the natural movie video stimulus (Figure 4**E**). We directly validated this hypothesis by evaluating the mutual information (MI) between the population spiking activity and the frames of the movie in a trial-by-trial manner in each state (Figure 4**G**, top). As expected, spiking activity in the high-frequency state was more informative about the stimulus than the lower-frequency state, with V1 neurons encoding most of that information (Figures 4**G**, bottom, S6**A**). In line with the observed high FF measures (Figure 4**F**), neurons in RL reported the lowest MI with the stimulus (see Discussion). Again, omitting the low MI measures in RL, pixel-level information decreased along the hierarchy during the high-frequency state (*S_H_* : Pearson’s r with anatomical hierarchy score = -0.90, p = 0.038; *SI* : Pearson’s r = -0.86, p = 0.06, *S_L_* : Pearson’s r = -0.81, p = 0.09). While these findings confirmed the association between spiking variability and stimulus representation across states, they further suggest a loss of pixel-level information along the visual pathway.

In summary, the high-frequency state is characterized by lower population shared variance, trial-to-trial variability, and increased spike timing variability (Figure S4**D-G**). During this state, variability trends showed strong anti-correlations with the anatomical hierarchy scores such that V1 demonstrated the highest variability across the different visual areas in all three measurements. This could be due to a strong influence of the temporal pattern of sensory drive in early areas, which is validated by the trend of decreasing pixel-level information encoded in V1, especially in the high-frequency state.

### HMM based predictor model

Given the substantial influence of the internal oscillation states on spiking variability and sensory processing, we next sought to quantify the impact of different variability sources on neural dynamics during the different states. We built an HMM-based linear encoding model to predict changes in single-trial neural activity in each visual area (Figure 5**A**). The resulting HMM-predictor model allows for the quantification of state-specific contributions of stimulus and other source variables to the target single-trial neural activity. Deriving inspiration from an HMM-GLM framework (Ashwood et al. 2022), the HMM-predictor model has two essential pieces: an HMM governing the distribution over latent LFP states (identified in the preceding section) and a set of state-specific predictors governing the weight distributions over the input features. However, unlike the previously proposed HMM-GLM, the state sequences are pre-determined by the HMM, and we do not re-train the HMM model for optimized prediction. Finally, the model also produces a time-varying kernel (*τ* seconds long) for each feature, relating that variable to neural activity in the subsequent time bin (Figure 5**A**, panel 3).

Our model considers an extensive array of variables that we classify into three categories: stimulus, behavior, and internal brain activity (Figure 5**A**, panel 1). Stimulus features include a set of higher (edges, kurtosis, energy, entropy) and lower-order (intensity, contrast) image features, and behavioral features include the complete set of movement variables determined in the previous section (see Figure 3). Under internal brain activity, the model includes both the averaged neuronal population activity from simultaneously recorded neighboring visual areas (that is, other than the target visual area) and the raw LFPs from different layers within the target area. Since model fits to linearly dependent input features are unreliable, we employed QR decomposition to systematically orthogonalize the input features (Mumford et al. 2015, see Methods).

We derived **two separate versions** of the HMM-predictor model to study neural variability at multiple scales: a population model and a single-neuron model (Figure 5**B**). The single-neuron model predicted the single-trial firing rate of the target neuron, while the population model predicted the single-trial averaged neuronal population activity in an area. In the population model, the predictors were linear regressors of the input features, and the model was fit using Ridge regression to prevent overfitting (equation 16). The single-neuron model accounted for the non-linearity associated with spike generation, wherein the predictors were designed as Poisson regressors of the input features, and the model was optimized by maximizing a regularized log-likelihood function to prevent overfitting (equation 17). To evaluate how well the model captured the target neural activity, we computed the five-fold cross-validated *R*^2^ (cv*R*^2^, equation 18).

### State specific contributions to population-level variability

The overall population model predicted 53.4 *±* 6.6% (mean *±* std, n = 25 sessions, Figure 5**C**) of the variance in the averaged neuronal population activity across the six visual areas. To evaluate the relative contributions from different source variables, we applied the model to individual sub-groups corresponding to each category. Interestingly, internal brain activity had the most predictive power (cv*R*^2^_1_ = 41.0 *±* 7.6%, mean *±* std, p = 2.5e-11, n = 25 mice), higher even than the combined power of behavioral and stimulus features (cv*R*^2^_*B+S*_ = 30.1 *±* 9.3%, mean *±* std, p = 0.0005, n = 25 mice). Stimulus features predicted the variance in the averaged neuronal population activity better than behavioral features (cv*R*^2^_*S*_ = 22.8 *±* 8.8%, cv*R*^2^_*B*_ 18.9 *±* 7.0%, mean *±* std, p = 0.009, n = 25 mice). These successive improvements in the explanatory power resulting from the inclusion of more sources are evident in the prediction traces shown in Figure 5**D**. It is worth noting that if single-neuron responses to external stimuli were completely independent, the contribution from stimulus features to population activity would be negligible. Nevertheless, the significant influence of stimulus features on population-level variability is suggestive of stimulus-related neuronal correlations within an area.

The addition of internal brain activity to the combined model of behavioral and stimulus features increased the explained variance by almost 24% (Δ*r*^2^_*F-(B+S)*_ = 23.5 *±* 10.2%, mean *±* std, 5**C**). Considering that LFP and population activity inherently carry information about stimulus and behavioral features, potentially making part of their contributions redundant, we have deliberately orthogonalized these internal variables against the stimulus and behavior variables (Mante et al. 2013). This orthogonalization ensures that internal variables capture variance beyond what can be accounted for by stimulus and behavior variables alone. To understand the substantial increase in explained variance, we analyzed the contributions of internal brain activity to each state. We found that these variables largely increased the predictability during the low-frequency state 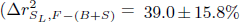, mean *±* std, Figure 5**E**, panel 1). Activity in this state was most poorly explained by the combined model of stimulus and behavioral features (cv*R*^2^_*S_L_,(B+S)*_ = 16.2 *±* 11.5%, mean *±* std, p = 8.3e-6, n = 25 mice). The combined model of stimulus and behavioral features was best at explaining variability in the high-frequency state, and accordingly, activity in this state showed a smaller improvement in its predictability on the inclusion of internal activity features 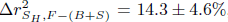, mean *±* std, p = 2.3e-6, n = 25 mice).

Consistently, within-area LFPs and averaged population activity from the neighboring visual areas contributed more towards explaining the activity in the low-frequency state (p = 4.5e-6, p = 8.2e-13, respectively; n = 25 mice, Figure 5**F**). At the same time, both stimulus and behavioral features demonstrated increased predictive power during the high-frequency state (p = 0.003, p = 0.01, respectively; n = 25 mice), suggesting a switch in the network dynamics within the visual cortex. The current findings are also consistent with prior studies (Lovett-Barron et al. 2017; McGinley et al. 2015; Speed et al. 2020), highlighting the role of slow-oscillatory waves in synchronizing spiking activity during the low-frequency state (Figure 4**C,D**), thereby disrupting stimulus encoding in this state.

**Figure 5:**
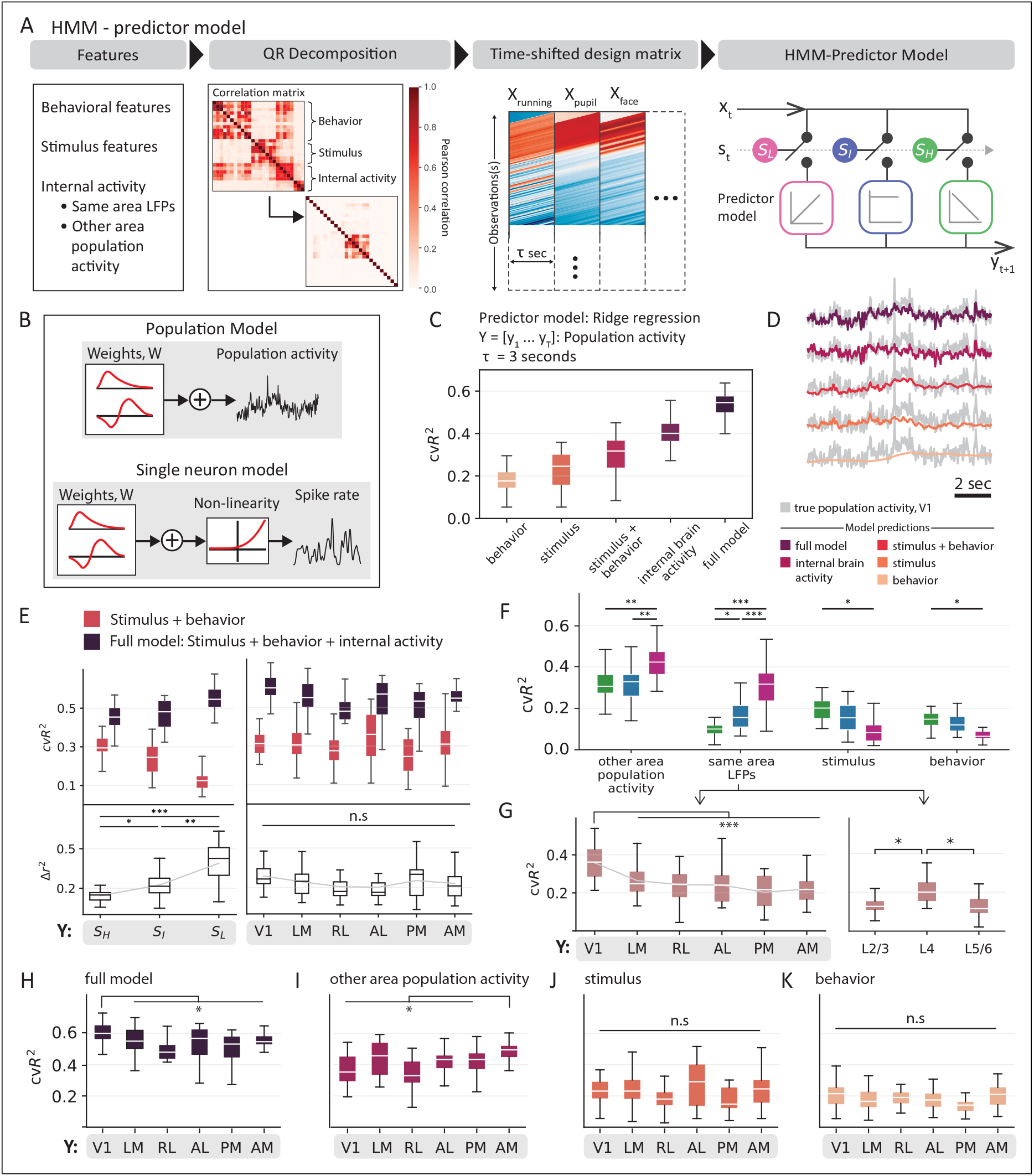
Relative contributions of the different sources to population-level variability. **A**, HMM-based prediction model to account for state-specific contributions of different sources of variability. Design matrices were constructed using decorrelated features to train state-specific regressors. *S_H_* : High-frequency state (green), *S_I_* : Intermediate state (blue), and *S_L_*: Low-frequency state (pink). **B**, HMM-predictor models to study encoding in population and single neuron models. Population models included a linear weighting of the input features, while in single neuron models, linear weighting was followed by a non-linear exponential projection. **C-G**, Results from population model. **C**, Explained variance for different categories of input feature groups, averaged across all mice obtained using five-fold cross-validation. The box shows the first and third quartiles, the inner line is the median over 25 mice, and the whiskers represent the minimum and maximum values. **D**, Averaged population responses overlaid with model predictions from respective input feature groups. **E**, Comparison of predictions in different (left) states and (right) visual areas prior to and post addition of internal brain activity. Top: Cross-validated explained variance for each model. Bottom: Unique contribution of internal brain activity. **F**, Contributions from single category models to explaining the variance in averaged neuronal population activity in different states. **G**, Contributions from LFPs in the same area to explain the variance in averaged neuronal population activity from (right) different layers and (left) in different visual areas. **H-K**, Same as G (left), but for different input features.

Using the complete set of input features, we could predict about 61.1 *±* 6.9% (mean *±* std, n = 25 mice) of the variance in V1’s averaged neuronal population activity, the highest among all visual areas (Figure 5**H**). Although including internal brain activity did not differentially affect predictability across visual areas (p = 0.12, n = 25 mice, Figure 5**E**, panel 2), contributions from its sub-components revealed interesting differences. Firstly, averaged population activity from neighboring areas explained more variance than within-area LFPs (p = 1.5e-9, n = 25 mice, Figure 5**F**). Secondly, their across-area prediction showed reversed trends. While LFPs explained significantly more variance in V1 than other visual areas (Figure 5**G**, panel 1), averaged population activity explained significantly more variance in AM (Figure 5**I**). Lastly, the predictive power of LFPs varied across the cortical depth, wherein layer 4 (L4) LFPs contributed more to the variance in the averaged neuronal population activity than LFPs in other layers (Figure 5**G**, panel 2).

When disregarding the influence of internal states, stimulus features did not significantly differ in their predictive power across areas (Figure 5**J**, p = 0.13), even at the level of single features (Figure S8**A, B**, p *∈* [0.33, 1]). However, state-specific analysis revealed pronounced differences in the high-frequency state (Figure S8**D, F**). In this state, different stimulus features also showed distinct predictive powers indicating heightened sensitivity to stimulus changes (Figure S8**C, E**). Specifically, higher-order stimulus features (edges, kurtosis, and energy) reported greater predictive power than stimulus contrast and intensity. Finally, facial movements made a more substantial contribution to the averaged neuronal population activity compared to other behavioral features (Figure S9**A, C**, p = 0.02, n = 25 mice), consistent with our observations in Figure 3**F**.

### State specific contributions to single-neuron variability

To explain the single-trial activity of individual neurons, we replaced the predictor in the HMM-predictor model with a GLM. This allowed us to systematically quantify the contributions from the different sources to single-neuron variability in each trial. Since a GLM predicts the conditional intensity of the spiking response, we evaluated our model performance against the rate functions of individual neurons obtained after smoothing the spike counts with a Gaussian filter (s.d. 50 ms). To appropriately identify their variability sources, neurons were further selected by a minimal firing rate (*>* 1 spikes/s in all states) and receptive field locations, along with the standard quality control metrics of the dataset (see Methods, Siegle, Jia, et al., 2021). After filtering, *n* = 3923 units remained across all mice and were analyzed using the GLM model.

Overall, the model was able to explain an average of cv*R*^2^_*F*_ = 26.7 *±* 13.5% (mean *±* std, *n* = 3923 units) of the total variance of single-trial activity across all neurons (Figure 6**A**) such that individual contributions from different sources showed a reversed trend compared to the population model. While the variance in the averaged neuronal population activity was best explained by internal brain activity, single neurons were best explained by stimulus and behavioral features [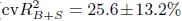, mean *±* std, *n* = 3923 units). Across all input features, stimulus features were most predictive of single-neuron activity (cv*R*^2^_S_ = 19.8 *±* 13.6%, mean *±* std), and LFPs were the least predictive (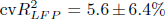, mean *±* std). However, the state-wise contribution trends of the individual input features were similar to that in the population model, such that neural activity was the most predictive during the high-frequency state (Figure 6**B**). Across areas, single neuron variability was best explained along the anterolateral path (LM, AM, and AL, cVR^2^_F_ = 26.2 ±0.9% (mean ± std), Figure S10, p = 2.5e-05). To aid visualization of the model predictions, we applied Rastermap (Stringer et al. 2019) to the spike counts of neurons, creating a 1-D embedding of the neural activity that captures their non-linear relations. Sorting the neurons by their eigenvalues revealed transient changes in the neural ensemble that were captured solely by the stimulus features (Figure 6**D**). Other features were less discerning and captured only the broad changes in the firing patterns.

Many features can impact an individual neuron’s variability, yet a specific feature often takes precedence. Accordingly, we grouped neurons based on the feature with the highest unique predictive power, explaining at least 10% of the unit’s spiking variance 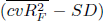. This categorization resulted in five distinct groups: one for each input feature and an additional group comprising neurons where no feature explained more than 10% of their variance. Examination of neuron distribution across visual areas revealed that the fraction of neurons best predicted by stimulus features peaked in V1, decreasing along the hierarchy (Figure 6**E**, Pearson correlation with hierarchy score, *r_p−RL_* = *−*0.91*, p_p−RL_* = 0.03). Conversely, fraction of behavior-related neurons increased along the hierarchy (*r_p_* = 0.89*, p_p_* = 0.04) such that *∼* 15 *−* 20% of neurons in areas between RL and AM were best predicted by behavior (RL: 21.5%, AL: 15.8%, PM: 15%, AM: 16.6%). Notably, despite the rise of behavior-related neurons in higher regions, the majority of neurons in each area were best explained by stimulus features (Figure 6**E**). Similar to behavior, number of neurons affected by neural activity from the neighboring areas also increased along the hierarchy (*r_p−RL_* = 0.95*, pp−RL* = 0.01). These findings indicate a rise in functional diversity among neurons ascending the visual hierarchy.

Next, we quantified variability based on the subgroups determined by the contributing features, employing metrics from the previous section. On average, neurons best explained by stimulus showed Fano factor (FF) values below 1, indicating sub-Poisson variability. These neurons also reported the lowest shared variance with other neurons in the population. In contrast, neurons primarily influenced by the averaged population activity from neighboring visual areas shared a large percentage of their variance with neurons in the target area, suggesting their involvement in internal synchronization. These neurons along with behavior-related neurons exhibited highest spike timing variability. It is important to emphasize that not all neurons encoded a single feature; 29% of neurons were well-predicted by multiple sources with cvR^2^ *>* 10% across all categories of input features: stimulus, behavior, and internal activity (Figure 6**C**). A thorough investigation of variability within this category of pan-modulated neurons would merit future research. Finally, the explained variance of fast spiking cells significantly surpassed that of regular spiking cells, except when stimulus features were used as input features, therefore, suggesting a greater involvement of regular spiking cells in stimulus encoding (Figure S10**F-K**, full model: p = 5.32e-11, behavior: p = 9e-48, internal activity: p = 0, stimulus: p = 0.05).

**Figure 6:**
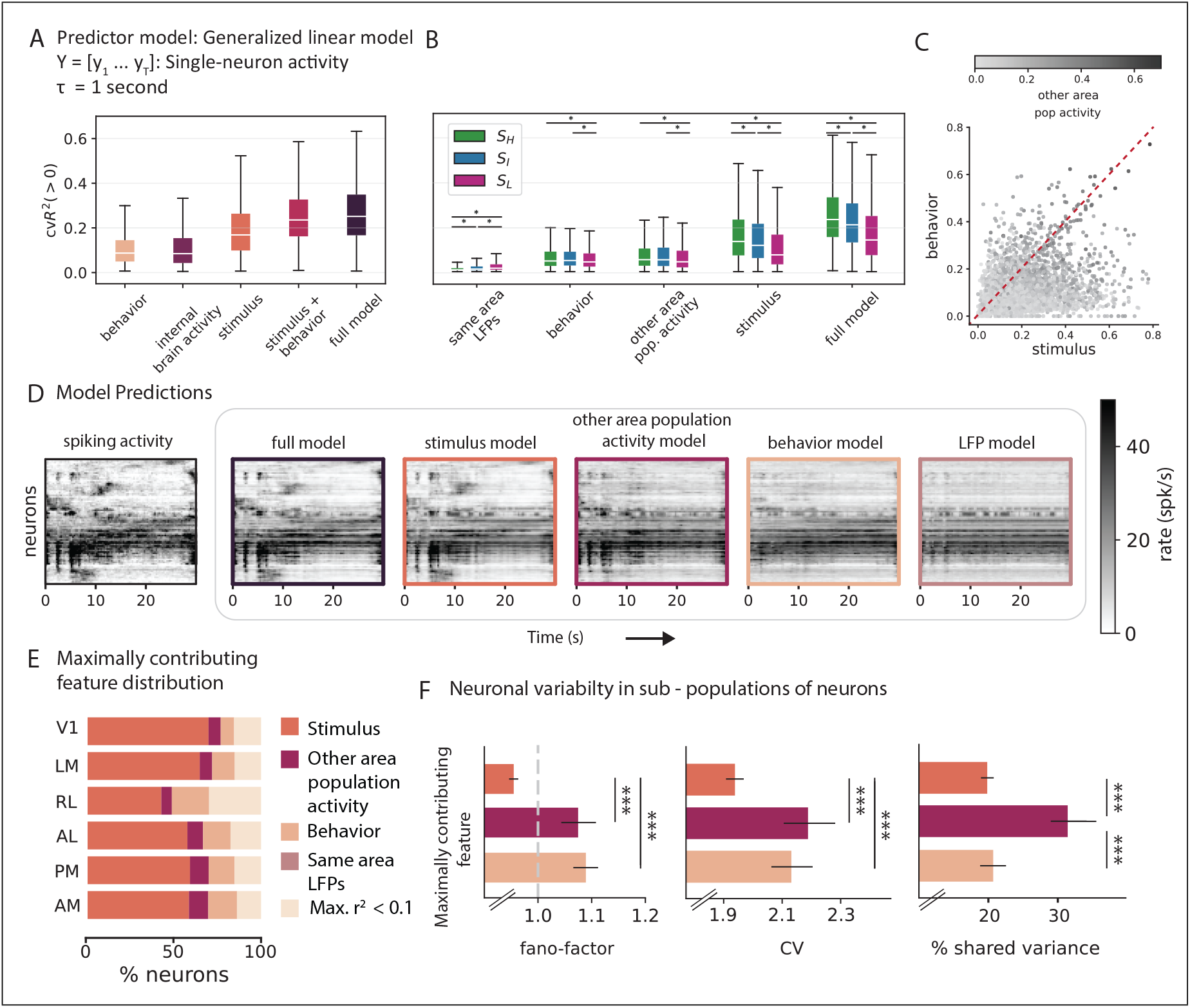
Relative contributions of the different sources to single neuron variability. **A**, Mean explained variance for different categories of input features, averaged across n = 3923 neurons and obtained using five-fold cross-validation. The box shows the first and third quartiles, the inner line is the median over all neurons, and the whiskers represent the minimum and maximum values. **B**, Contributions from single category models to explaining single-neuron variability during different oscillation states. *S_H_* : High-frequency state (green), *S_I_* : Intermediate state (blue), and *S_L_*: Low-frequency state (pink). **C**, Explained variance of all units in each input feature category. **D**, (First panel) Neuronal activity, with neurons sorted vertically by a manifold embedding algorithm, Rastermap. (Panels 2 - 6) Prediction of neuronal activity (n = 350 units, best explained units across mice and areas) from respective input feature categories. **E**, Proportion of units in each area with maximal explained variance from respective input feature categories. No units were maximally explained by LFPs from the same area. **F**, (left to right) Variability across trials (Fano factor), variability across time (coefficient of variation), and shared variability of neurons grouped according to their maximally contributing feature.

## 3 Discussion

Our observations provide a comprehensive description of the non-stationary aspects of spiking variability in the visual cortex as the brain traverses through distinct oscillation states. We characterized this variability along three dimensions: variability across trials (Kara et al. 2000), variability in spike times (Softky and Koch 1993), and shared variance within a population (Williamson et al. 2016). By utilizing cortical LFPs to define different internal oscillation states, we found that each state captured a distinct profile of spiking variability. Using the state fluctuations as a temporal backbone, we incorporated the non-stationary properties of neuronal variability into an HMM-based encoding model. The linear encoding model was able to partition and evaluate the relative contributions from three different sources of variability: visual stimulus, behavior, and internal brain dynamics, explaining single-neuron variability with 27% and averaged population activity with 53% accuracy. Each source influenced spiking variability in a state and area-specific manner. Overall, our study not only underscores the importance of addressing the non-stationary dynamics of spiking variability, but also emphasizes the imperative to account for the dynamic influence of the internal and external factors on stimulus representation (Figure 7).

### Relative influence of different sources on neuronal variability

Identifying and locating the different sources influencing neural variability poses a significant challenge in systems neuroscience (Goris et al. 2014; Renart and Machens 2014). Previous research has emphasized the significance of internal brain activity in accounting for neuronal variability (Carandini 2004; Schöolvinck et al. 2015; Shadlen and Newsome 1998). While these studies did not consider variability induced by externally observable task- and behavior-related variables, recent investigations have predominantly focused on this latter category of input features (Musall et al. 2019; Recanatesi et al. 2022; Steinmetz et al. 2019; Stringer et al. 2019). In this study, we adopt a comprehensive approach by integrating contributions from both internal brain activity and externally observable variables to understand neuronal variability.

We considered a two-fold contribution from internal brain activity. Firstly, utilizing brain states defined by internal oscillatory rhythms as a temporal framework, we were able to associate the various dynamics of spiking variability with these internal states. Secondly, we incorporated averaged neuronal population activity from each neighboring area and LFPs as input features into the HMM-based encoding model. These variables played a significant role in explaining neural variability, primarily contributing to activity in the low-frequency state. Consistent with previous findings (Carandini 2004; Schöolvinck et al. 2015), internal variables explained approximately 40% of the total variability of averaged neuronal population activity within an area, even surpassing the variance explained by the combined model of stimulus and behavioral features by 11% 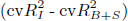 At the level of single neurons, contributions from internal brain activity, although relatively small, remained statistically significant, explaining around 11% of the total variance. However, this was nearly 14% 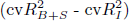 less than the variance explained by the combined model of stimulus and behavioral features.

Recent progress in behavioral video analysis, computational modeling, and large-scale recording techniques has highlighted the impact of movement-related variables on neural activity across the cortex (Musall et al. 2019; Steinmetz et al. 2019; Stringer et al. 2019). Our observations are consistent with these findings. Behavior-related variables explained up to *∼* 20% of the averaged neuronal population activity and *∼* 12% of single-neuron variability in the visual cortex. Moreover, the influence of behavior becomes more pronounced in the high-frequency state (Figures 5**C**,6**A**, 7) and as one ascends the visual hierarchy, entraining a larger proportion of the neural population (Figure 6**E**). However, our findings diverge from those reported in Musall et al. 2019, which found that uninstructed movements exerted a greater influence on V1 neural activity than a visual stimulus. We attribute this difference to three reasons: first, our mice are passively viewing the screen without engaging in a behavioral task; second, our naturalistic movie stimulus may engage a broader array of neurons compared to the static, flashed stimuli used in previous research; third, our recording captures single-unit spiking activity, contrasting with previous wide-field calcium imaging. In addition to behavior, these differences underscore the importance of recording methodologies, experimental conditions and stimuli, prompting a closer examination of the specific factors influencing single-trial neural activity in diverse contexts.

Despite large variability in spiking activity, neuronal populations exhibit a remarkable ability to robustly encode information across different brain regions (Harris et al. 2019; Jia et al. 2022; Perkel and Bullock 1968). Our results suggest this is state-dependent. A clear pattern emerges throughout our analyses: population dynamics during the high-frequency state are the most effective in representing stimulus information, while stimulus features weakly modulate activity in other states ((Figures 4**G**, 5**F**, 6**B**), 7). While several lines of studies have indirectly confirmed this state-dependence of information encoding either through reports of task performance or via investigations under artificially induced states of anesthesia (Haider et al. 2007; Mccormick et al. 2020; Poulet and Petersen 2008; Scholvinck et al. 2015), our findings directly quantify and describe this dependency. Specifically, we find that spiking activity in the high-frequency state has the lowest shared variance, lowest trial-to-trial variability, and the highest spike timing variability (Figure 4). These characteristics of single-neuron activity may result from enhanced encoding of various temporal and spatial features of the time-varying natural movie stimulus during the high-arousal state (Figures 5**F**, 6**B**). In contrast, the dominance of slow oscillatory activity in low-frequency state, coupled with high shared variance, trial-to-trial variability, and more regular firing, appears to reflect internal dynamics that disrupt the accurate representation of stimulus information. We posit that this observed correlation between heightened sensory encoding capacity and increased arousal during the high-frequency state may arise from the mice’s innate survival mechanism, leading them to enhance visual information intake while in a state of heightened alertness or running.

**Figure 7:**
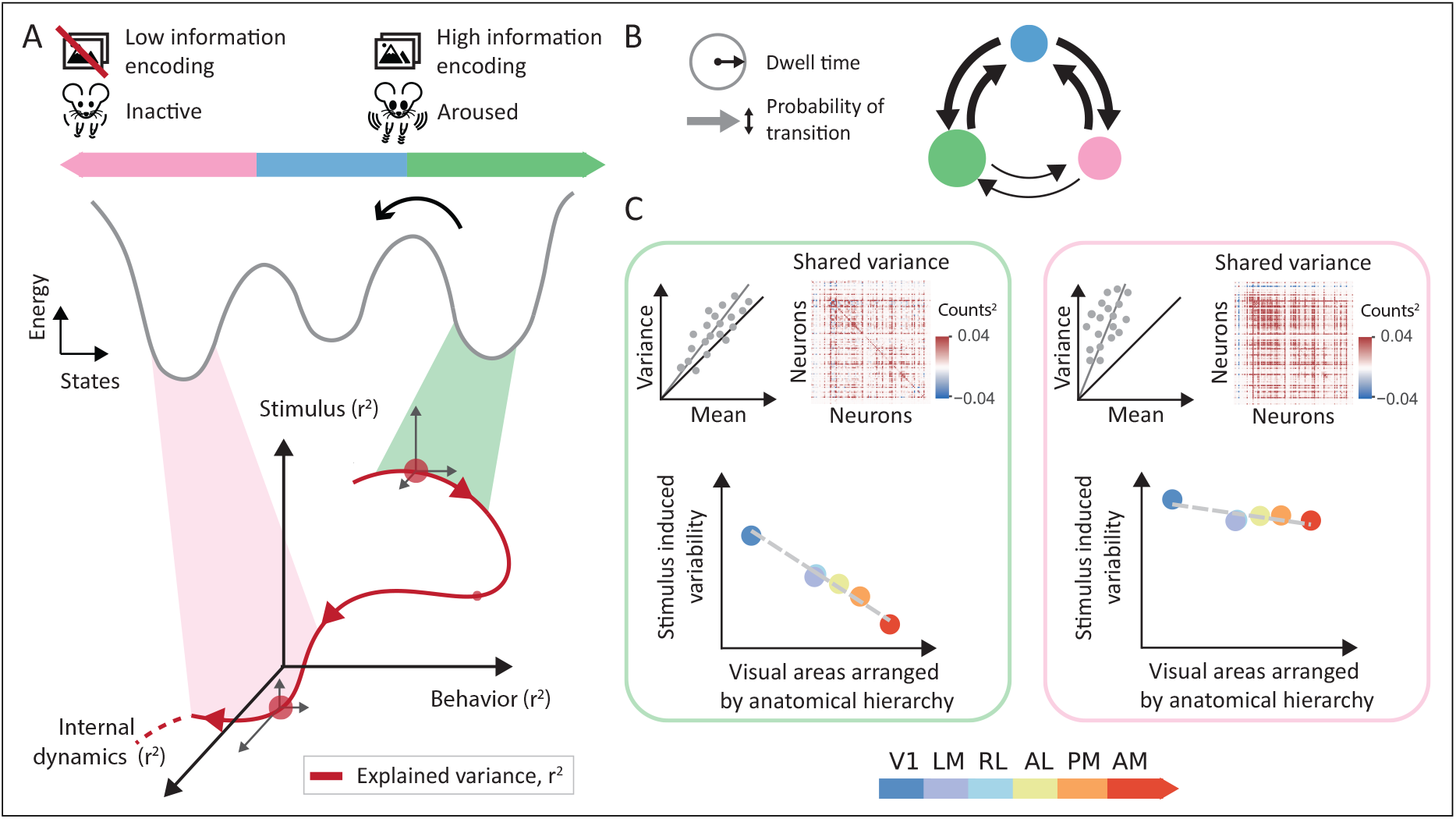
Illustration of the dynamic nature of neuronal variability and sensory encoding. **A** Illustration of variance explained (red line) in neural activity by stimulus (z-axis), behavior (y-axis) and internal dynamics (x-axis), along with their respective associations with internal state, as delineated by the basins of attraction in the energy landscape of neural activity (gray line). The interplay between internal and external factors collectively shape the time course of neuronal variability, influencing sensory coding. States serve as a temporal framework underlying the dynamic nature of these interactions. **B** Graphical depiction of state transitions. Transitions between low and high-arousal states occur via an intermediate state, with the least amount of time spent in this intermediary phase. In mice, each state typically only lasts for a short duration of 1-2 seconds. Transition probabilities are depicted by the thickness of arrows, while the duration of each state is indicated by the size of the circles. **C** Illustration of state-specific profiles of spiking variability (Fano factor and shared variance). Left panel: Neurons in the high arousal state demonstrate improved stimulus encoding characterized by lower trial-to-trial variability and lower shared variability within a population. In this state, stimulus induced variability gradually decreases along the visual hierarchy (Harris et al. 2019). Right panel: In low arousal states, stimulus effects are hindered by internal dynamics that predominantly influence the observed neuronal variability. Neuronal activity in this state is highly synchronous within and across areas and demonstrates higher trial-to-trial variability.

### Sensory processing along the visual cortical hierarchy

Given the hierarchical organization of the visual cortex (Siegle, Jia, et al., 2021,Harris et al. 2019), the response variance of a sensory neuron can potentially limit the amount of stimulus information available to downstream circuits (Denman and Reid 2019, Figures 4**G**, 7). While past studies have shown the effects of pair-wise correlations on information encoded by a neuronal population (Averbeck et al. 2006; Kohn et al. 2016; Moreno-Bote et al. 2014), a more comprehensive population-level perspective is essential to understanding the brain’s correlational structure (Recanatesi et al. 2022; Shea-Brown et al. 2008; Trousdale et al. 2012). Here, we applied shared variance (Williamson et al. 2016) as a generalization of the pair-wise correlations between single neurons extended to an entire population. Notably, we observed a decrease in the percent of shared variance along the visual hierarchy (Figure 4**D**). While this decline might imply the introduction of independent noise at subsequent stages of signal processing, it could alternatively result from the increased diversity of neurons influenced by factors other than the stimulus itself (Figure 6**F**). The high variance shared across neurons in V1 can likely be attributed to V1 comprising the largest proportion of neurons exhibiting strong, time-locked responses to the temporal dynamics of stimulus features (Figure 6**F**, Churchland et al. 2010; Matteucci et al. 2019). Our findings provide further support for this notion, particularly through the observation that neurons in V1 reported high spike-timing variability, likely corresponding to the variance induced by a constantly changing stimulus(Figure 4**E**). Consistently, LFPs have a more pronounced influence on averaged population activity in V1 in comparison to other visual areas (Figure 5**G**). This suggests that the collective synaptic inputs into V1, represented by LFPs in the area, may entrain a larger population in V1 than in other areas.

Previous studies have indicated that trial-to-trial variability (Fano factor) increases as information propagates up along the visual pathway from the retinal receptors to the primary visual cortex (Bair 1999; Kara et al. 2000; Schöolvinck et al. 2015). Our observations mirror this trend in the visual cortex when mice were exposed to full-field light flashes, revealing an increase in trial-to-trial variability along the cortical hierarchy (Figure S6**D**). However, in response to natural movies, trial-to-trial variability decreased along the visual cortical hierarchy (Figure 4**F**). We attribute this decrease in variability to the heterogeneous properties of a natural movie frame where, in awake mice, eye movements (even small saccades) across the frame could elicit more variable neuronal responses across trials in early visual areas with smaller receptive fields (Gur et al. 1997). Lastly, it is important to note the variability properties of neurons in the rostrolateral visual area (RL), which do not always follow the visual hierarchy trends. This is especially true when considering trends related to stimulus encoding, such as trial-to-trial variability and mutual information (Figures 4**F,G**, 6**E**, S6**C, D**). We attribute this to two reasons. Firstly, since RL is located at the border of the visual and primary somatosensory (S1) cortices, the functional specialization of neurons in RL is likely more diverse than in other visual areas. This is reflected in our findings where RL had the smallest proportion of neurons influenced by stimulus features and the largest proportion of neurons with low explained variance (Figure 6**F**). Secondly, due to the retinotopic center of RL being situated on the boundary between RL and S1 (Olcese et al. 2013), it is often challenging to target its precise retinotopic center (de Vries et al. 2020).

### Dynamic shifts in neuronal variability

The dynamic nature of neuronal variance across time has been consistently demonstrated in theoretical and empirical analyses (Churchland et al. 2011; Goris et al. 2014; Stein 1965). Here, we specifically quantify the magnitude of stimulus-driven neuronal variability associated with internal states. Our findings show that, during passive viewing, mice typically persist in a specific state for an average duration of 1.5 *±* 0.1 seconds, indicating that state-dependent neuronal variability undergoes changes within seconds (Figures 2**F**, 7). The state sequences reveal a smooth transition of neuronal variability between distinct variability profiles, passing through an intermediate state (Figures 2**G**, 4, 7). Moreover, each state constitutes a unique composition of sources that influence neuronal variability (Figures 5**F**, 6**B**). These rapid shifts in source composition across states arise from the complex interactions between non-stationary source variables, collectively contributing to the non-stationarity of neuronal variability (Figure 7).

These findings offer additional insights into the dynamic properties of neuronal variability, providing important constraints for theoretical modeling of stimulus-driven variability. Firstly, the dynamically changing source composition indicates that the responsiveness of a neuronal population to sensory input varies over time, challenging the assumption of a constant stimulus contributing to the responsiveness of a sensory system. Secondly, accounting for the distinct variability profiles associated with different internal states can specifically address the non-stationary stimulus-encoding capability of neuronal populations. Lastly, integrating state fluctuations as a temporal framework can enhance our understanding of the network dynamics contributing to non-stationary neuronal variability.

### Future directions

Considering the differences in stimulus representation across states, we expect these states to similarly influence the accurate transmission of sensory-related information. Interestingly, it has been shown that artificially inducing synchronized low-frequency oscillations in area V4 of the primate visual cortex impairs the animal’s ability to make fine sensory discriminations (Nandy et al. 2019). Studies in mice have also found that slow-oscillatory activity in key-sensory areas, such as the somatosensory, visual, and auditory cortex, significantly reduces their ability to quickly and accurately respond to sensory stimuli (Bennett et al. 2013; Crochet and Petersen 2006; McGinley et al. 2015). These studies suggest a disruptive impact of the local slow oscillatory activity on downstream cortical processing. Our current findings indicate that during the low-frequency state, reduced stimulus influence on spiking activity diminishes area-wise differences in variability along the visual hierarchy (Figures 4, 7). While shared variance and trial-to-trial variability increase in the low-frequency state, their trend across the hierarchy flattens in this state, suggesting a lack of differentiation in how these regions respond to sensory-related information. To understand these effects, our future investigations will focus on network properties and information propagation as the brain transitions through the various oscillation states.

In this study, we make use of the controlled yet dynamic structure of the passive viewing design to trace neuronal variability across discrete oscillation states in awake mice. While our discrete characterization of brain states provides a straightforward interpretation of neural activity, recognizing the possibility of continuous state changes (such as a continuum of pupil size or network activity changes) is vital for exploring the full spectrum of neural responses in awake, behaving animals. Additionally, to fully characterize neuronal variability and its influence on information processing in the cortex, investigating neural activity during active tasks is essential. Recent studies have shown that a subject’s engagement during an active task varies drastically from trial to trial, playing out through multiple interleaved strategies (Ashwood et al. 2022; Piet et al. 2023; Zhuang et al. 2021). While the tools in this study can help identify variables that promote task engagement, they do not elucidate the underlying mechanisms causing state transitions. Understanding these dynamics entails a thorough investigation of unit activity in the subcortical and dopaminergic regions of the brain.

Our observations, combined with existing studies on spiking variability, suggest that cortical state acts as a key determinant of the variability seen in the cortex. By offering a comprehensive view of this variability, we have been able to directly study both the sensory and non-sensory aspects of neuronal responses in the visual cortex. It is evident that spiking variability in the cortex transcends mere ‘neural noise’, and explaining neuronal variability by partitioning it into different origins can help us understand its influence on information representation and propagation in the brain, and ultimately resolve its computational contribution to behavior.

## 4 Methods

### Data Collection

The data analyzed and discussed in this paper are part of the publicly released Allen Institute Brain Observatory Neuropixels dataset (n=25 mice) (Siegle, Jia, et al., 2021). Neural recordings used Neuropixels probes (Jun et al. 2017) comprising 960 recording sites. Either 374 for “Neuropixels 3a” or 383 for “Neuropixels 1.0” were configured for recording. The electrode sites closest to the tip formed a checkerboard pattern on a 70 *µ*m wide x 10 mm long shank. Six Neuropixels probes were inserted at the shallowest 2 mm and at the deepest 3.5 mm into the brain for each recording. These requirements ensured adequate recordings of the cortex while preventing any brain damage. To ensure that the probes were recording from functionally related cells in each visual area, retinotopic centers were determined and targeted accordingly. Targeting the cortical visual areas, AM, PM, V1, LM, AL, and RL, was guided by the angle of approach of the probe, as well as the depth of functionality of the imaging boundaries. All procedures were performed according to protocols approved by the Allen Institute Institutional Animal Care and Use Committee under an assurance with the NIH Office of Laboratory Animal Welfare.

The Open Ephys GUI was used to collect all electrophysiological data. Signals from each recording site were split into a spike band (30 kHz sampling rate, 500 Hz highpass filter) and an LFP band (2.5 kHz sampling rate, 1000 Hz lowpass filter). Spike sorting followed the methods outlined in Jia et al. 2022. Briefly, the spike-band data was subject to DC offset removal, median subtraction, filtering, and whitening before applying the Kilosort2 MATLAB package (https://github.com/MouseLand/Kilosort) for spike time identification and unit assignment (Stringer et al. 2019). Detailed information about the complete experimental design can be found in Durand et al. 2022.

### Statistics and data analyses

For all analyses, Python was used as the primary programming language. Essential analytical tools utilized include Scipy (Virtanen et al. 2020) and Scikit-learn (Pedregosa et al. 2011). Error bars, unless otherwise specified, were determined as the standard error of the mean. For comparisons across units (n = 7609 units after QC filtering, and n = 3923 units post-RF filtering), mice (n = 25), or states (n = 3), we used a one-way ANOVA for Gaussian-distributed metrics and the rank sum test for non-Gaussian distributed metrics. In cases of high subject-to-subject variability, we used a paired t-test. Bonferroni correction was applied for multi-group comparisons. To determine if a distribution significantly differs from zero, we used a one-sample t-test. To evaluate the similarity to the previously established anatomical visual hierarchy in mice (Harris et al. 2019), we computed the correlation between our measured variable and the anatomical hierarchy score (V1: -0.50, RL: -0.14, LM: -0.13, AL: 0.00, PM: 0.12, AM: 0.29), and Pearson’s correlation was applied to estimate the significance of correlation.

### Visual Stimulus

Custom scripts based on PsychoPy (Peirce, 2007) were used to create visual stimuli, which were then presented on an ASUS PA248Q LCD monitor. The monitor had a resolution of 1920 x 1200 pixels and a refresh rate of 60 Hz, measuring 21.93 inches wide. The stimuli were shown monocularly, with the monitor positioned 15 cm from the right eye of the mouse. The visual space covered by the stimuli was 120*^◦^* 95*^◦^* before any distortion occurred. Each monitor used in the experiment was gamma corrected and maintained a mean luminance of 50 cd/m^2^. To accommodate the mouse’s close viewing angle, spherical warping was applied to all stimuli to ensure consistent apparent size, speed, and spatial frequency across the monitor from the mouse’s perspective.

#### Receptive field mapping

The receptive field locations were mapped with small Gabor patches randomly flashed at one of 81 locations across the screen. Every degree of drifting grating (3 directions: 0*^◦^*, 45*^◦^*, 90*^◦^*) was characterized by a 2 Hz, 0.04 cycles with a 20*^◦^* circular mask. The receptive field map (RF) for an individual unit is defined as the average 2D histogram of spike counts at each of the 81 locations, where each pixel corresponds to a 10*^◦^ ×* 10*^◦^* square.

#### Stimuli for passive viewing

The mice were exposed to various types of stimuli during the experiment, including drifting gratings, natural movies, and a flashes stimulus. The gratings stimulus included 4 directional gratings that were repeated 75 times at a frequency of 2 Hz. As for the natural movies, they were divided into 30-second clips, and each clip was repeated 30 times as a block. To introduce variability, there were an additional 20 repeats with temporal shuffling. Lastly, the flashes stimulus included a series of dark or light full field image with luminance = 100*cd/m*^2^.

### Quality control metrics

All single-neuron analyses (Figures 4, 6) were performed on neurons that successfully met three essential quality control thresholds: presence ratio (*>* 0.9), inter-spike interval violations (*<* 0.5) and amplitude cut-off (*<* 0.1). Specific details of these metrics can be found in (Siegle, Jia, et al., 2021). These metrics were implemented to prevent the inclusion of neurons with noisy data in the reported analyses, considering both the physical characteristics of the units’ waveforms and potential spike sorting challenges. For single-neurons analyzed in Figure 6, a tighter threshold on presence ratio (*>* 0.95) was incorporated to avoid inflated values of prediction accuracy. Additionally, analyses in Figures 4**F** and 6 were filtered for neurons with receptive fields positioned at least 20 degrees away from the monitor’s edge. This criterion was incorporated to facilitate a meaningful comparison of the relative contributions from different sources of variability.

### Local field potentials and time-frequency analysis

Prior to constructing the hidden Markov model (HMM), we identified appropriate frequency ranges in the LFPs. To evaluate their power spectra, we applied short time-Fourier transform (STFT) on single channels using a Hann window of size *∼* 800 ms such that consecutive windows overlapped over *∼* 400 ms. Z-scoring the power spectrum at each frequency revealed LFP modulations in distinct frequency bands (Figure 2**B**). Further informed by the literature on LFPs in the mouse cortex (Akella et al. 2021; Buzśaki and Draguhn 2004; Fries 2015; Jia and Kohn 2011; Lundqvist et al. 2016), the following frequency ranges were selected from the LFP spectrum: 3-8 Hz (theta), 10-30 Hz (beta), 30-50 Hz (low gamma), and 50 - 80 Hz (high gamma). To filter the LFPs, we constructed four IIR Butterworth filters of order 11, each corresponding to the above frequency ranges. Finally, envelopes of the filtered LFP signals, obtained via the Hilbert transform, were supplied as inputs to the HMM.

The input features of the HMM model incorporate LFPs from across the cortical depth. To determine the corresponding layer of each LFP channel, we first estimated the depth of the middle layer of the cortical column. Similar to methods summarized in Stoelzel et al. 2009 and Jia et al. 2022, we applied current source density (CSD) on the LFPs within the 250 ms interval post-presentation of the flashing stimulus. To evaluate the CSD, we calculated each recording site’s average evoked (stimulus-locked) LFP response (*s*) and duplicated the uppermost and lowermost LFP traces. Next, we smoothed the signals across sites as shown in equation 1, where *r* is the coordinate perpendicular to the layers, and *h* is the spatial sampling distance along the electrode. Finally, the CSD mapping was obtained as the second spatial derivative of the LFP response (equation 2, Figure S1**D**, right). The CSD map can approximately dissociate the current sinks from current sources, respectively indicated as downward and upward deflections in the density map.

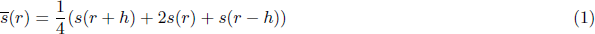

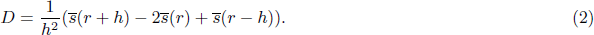

To facilitate visualization, we used 2D Gaussian kernels (*σ_x_* = 1*, σ_y_* = 2) to smooth the CSD maps. We identified the location of the input layer based on the first appearance of a sink within 100 ms of the stimulus onset. We then designated the center channel of the middle layer (L4) as the input layer and marked eight channels above and below it as L4. All channels above the middle layer were classified as superficial layers (L2/3), while all channels below the middle layer but above the white matter were categorized as deep layers (L5/6). Lastly, for each mouse, we validated the layer classification against the spectral decomposition of the LFPs across depth (Figure S1**D**).

### Identification of internal oscillation states - Hidden Markov model

We used a hidden Markov model (HMM) to detect latent states or patterns from envelopes of band-passed LFP signals. According to the model, network activity along the visual hierarchy is in one of *M* hidden “states” at each given time. Each state is a vector, *S*_(*a,d*)_, constituting a unique LFP power distribution over all depths (d = [L2/3, L4, L5/6]) across six visual areas (*a* = [*V* 1*−AM*]) in the cortex (emission matrix, Figure S3**A**). In an HMM-based system, stochastic transitions between states are assumed to behave as a Markov process such that the transition to a subsequent state solely depends on the current state. These transitions are governed by a “transition” probability matrix, *T_m,n_*, whose elements represent the probability of transitioning from state m to state n at each given time (Figure S3**B**). We assumed the emission distribution to be a Gaussian distribution over the power signals to train a single HMM for each mouse, yielding the emission and transition probabilities between states. To match the frame rate of the natural movie, we averaged the power signals within non-overlapping windows of 30 ms. Each HMM was optimized using the Baum-Welch algorithm with a fixed number of hidden states, M.

In an HMM, the number of states, M, is a hyperparameter. To find the optimum number of states (*M ^∗^*) per HMM, we optimized the 3-fold cross-validated log-likelihood estimate, penalizing the metric if the inferred latent states were similar. The correction for similarity was imperative to determining distinct states with unique definitions. ‘Similarity’ between the states was quantified as the top eigenvalue of the state definition matrix evaluated as the mean power across the identified frequency ranges (number of states *×* number of frequency bands, Figure 2**C**, right). The top eigenvalue represents the maximum variance in the matrix. In such a case, smaller values indicate lower variance in the definition matrix and, therefore, highly collinear state definitions. To apply this correction, we divided the log-likelihood estimate with the top eigenvalue where both metrics were individually normalized between -1 and 1 over a range of M *∈* [2, 6]. Normalization was performed to allow equal weighting of the two metrics. The log-likelihood estimate increases with the number of states until reaching a plateau, while the value of the top eigenvalue decreases. A ratio between the two metrics consistently pointed to *M ^∗^* = 3 optimal states across all mice (Figure 2**C**).

To further validate our model selection, we used the k-means algorithm as a control to cluster all the input LFP variables between *k* = 2 and *k* = 6 clusters (Figure S2**A**). To determine the number of clusters (states), we applied the Elbow method to the percentage of variance explained by each clustering model. The percentage of explained variance is the ratio of the variance of the between-cluster sum of squares to the variance in the total sum of squares. Applying the elbow method to each mouse, we selected the number of clusters, *k^∗^*, for which the incremental increase in the explained variance had the largest drop (the point of largest curvature) before the plateau (Satopaa et al. 2011). In most mice, the LFPs optimally clustered into three or four separate groups, displaying a remarkably similar power distribution obtained via the HMM. As a final sanity check, we applied dimensionality reduction to the input LFP variables using UMAP (Uniform Manifold Approximation and Projection, McInnes et al. 2018) and evaluated the silhouette scores (*sklearn.metrics.silhouettescore*) on the reduced input matrix based on the HMM states. The distribution of the silhouette scores across all mice further confirmed our model selection (Figure S2**B**).

The input LFP variables supplied to the HMM model include LFPs from one randomly selected channel from each layer of the cortical column: L2/3, L4, and L5/6, across all six visual area. This approach aims to achieve smoother states by reducing the number of input variables provided to the HMM model while ensuring representation across the cortex. We validated this input selection using two controls. First, we tested if latent states varied across visual areas. For this, we estimated HMM states using LFPs from each individual area (Figure S1**B**). Second, we conducted a randomized control test for each session, running 20 independent HMM fits with randomly selected LFP channels from each layer (Figure S1**E**). The initial guesses for emissions and transition probabilities were kept constant across different runs. Subsequently, for each test, we evaluated the pairwise correlations between state predictions for each pair of the HMM models. The correlation coefficients averaged around 0.54 *±* 0.04 (mean *±* sem, n = 25 mice, Figure S1**C**) for the area-wise control and around 0.75 *±* 0.04 (mean *±* sem, n = 25 mice, Figure S1**F**) for the layer-wise control, indicating the robustness of the determined states against area and channel selection.

### Behavioral features

Two synchronized cameras were used to record the mice: one focused on the body at a 30 Hz sampling rate, and the other an infrared camera focused on the pupil at a 60 Hz sampling rate. Running wheels were equipped with encoders to measure distance and speed of the mouses’ running during the data acquisition session. Behavioral variables used in regression analyses were quantified using universal mouse models constructed using DeepLabCut (Nath et al. 2019, Siegle, Jia, et al., 2021) for pupil size changes and using SLEAP (Pereira et al. 2022) for limb-to-tail movements. SLEAP, a modular UNet-based machine learning system, was trained to recognize up to 7 tracking points on the mouse’s body, including the body center, forelimbs, hindlimbs, and the proximal and distal ends of the tail (Figure 2**C**). However, the right forelimb was frequently occluded from view and subsequently dropped from our analyses. We trained the model on a combined 1311 labeled frames from across all mice, with annotations ranging from 10 to 300 frames per mouse. Utilizing SLEAP’s human-in-the-loop workflow, we alternated between labeling and training the model to achieve incremental improvements in prediction. In frames with resolutions of 478 *×* 638 pixels, the final model reported an average pixel error of 7.15 *±* 4.1 (mean *±* std, n = 1311 frames) pixels across all body parts. Input features for the regression models were generated as smoothed Euclidean distances between coordinates of each body part in consecutive frames. Additionally, facial movements were quantified using face motion energy from cropped behavior videos (Stringer et al. 2019). At each time point, this energy was determined as the sum of the absolute differences between consecutive frames. Lastly, the full set of methodological details for pupil tracking can be found in Siegle, Jia, et al., 2021.

### Variability metrics

#### Shared Variance

To investigate the co-variation of diverse neurons within a population, we employed linear dimensionality reduction techniques, as summarized in (Williamson et al. 2016). Specifically, we utilized factor analyses (FA) to quantify the percentage of variance shared across neural populations in the visual cortex. FA explicitly divides the spike count covariance into two components: a shared component and an independent component. The shared component captures the variability that is common across neurons within the recorded population, while the independent component quantifies the Poisson-like variability specific to each individual neuron. The FA analysis is performed on a matrix, 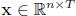 comprising spike counts from *n* simultaneously recorded neurons, along with a corresponding mean spike count vector, 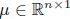. As illustrated in Figure 4**D**, FA effectively separates the spike count covariance into the shared component represented by *LL^T^* and the independent component represented by Ψ.

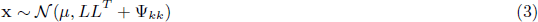

Here, *L ∈* R*^n×m^* is the loading matrix that relates the ‘*m*’ latent variables to the neural activity, and Ψ is a diagonal matrix comprising independent variances of each neuron. We calculated the percent shared variance for each neuron by utilizing the model estimates of the loading matrix, *L*, and the diagonal matrix, Ψ. This enabled us to quantify the degree to which the variability of each neuron was shared with at least one other neuron within the recorded population. For the *k^th^* neuron, the percent shared variance was evaluated as follows:

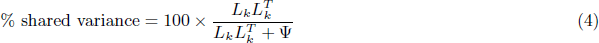

For our analyses, the FA model parameters, *µ, L*, and Ψ, were estimated using singular-value decomposition (sklearn.decomposition.FactorAnalysis). The number of latent variables, *m*, was determined by applying FA to the spike counts and selecting the value for *m* that maximized a three-fold cross-validated data likelihood (m = 24 *±* 3 factors, mean *±* std). Spike counts were evaluated in 30 ms bins and all values of shared variance reported in the paper (Figures 4**D**, 6**F**) present averages over all neurons in the given analyses. In Figure 4, state-specific shared variance for each neuron was evaluated on spike count matrices, **x** *∈* R*^n×Ts^*, comprised of concatenated epochs from each state. This allowed us to assess how much variability each neuron shared with others during specific oscillation states.

#### Coefficient of Variation

In our study, we investigate the spike timing variability of single neurons by analyzing the distributions of their inter-spike-intervals (ISIs). To achieve this, we constructed histograms of the ISIs and quantified their characteristics using the coefficient of variation (CV). The CV is a dimensionless metric that represents the relative width of the ISI histogram. It is calculated as the ratio between the standard deviation of the ISIs (*σ*_Δ_*_t_*) and their mean 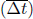.

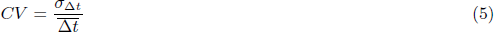

To evaluate the coefficient of variation (CV) of individual neurons in different states, we created histograms of their inter-spike-intervals (ISIs) based on the spike times observed within each state. However, since the large differences in the dwell times of different states would bias the range of ISIs in each state, and, consequently, the state-specific coefficients, we fixed the range of the ISI histograms. We chose an interval of t = 2.5s, the range at which incremental increase in CV had the largest increase (the point of largest curvature) before it plateaued (Figure S6**B**). Finally, values of CV reported throughout the paper (Figures 4**E**, 6**F**), represent the average across all single neurons in the given analyses.

#### Fano Factor

We evaluated the trial-to-trial variability of neuronal activity in the visual cortex using Fano factor (FF), calculated as the ratio of variance to the mean spike count across trials, respectively. Similar to previous studies in the visual cortex (Kara et al. 2000; Softky and Koch 1993), we computed the FF of each neuron within non-overlapping windows of 150 ms and averaged it across time. However, quantifying trial-wise variability of single neurons in a state-specific manner posed challenges. Partitioning each session into states over time disrupted the trial structure, necessitating an additional constraint over the number of trials in each window. A time-window was considered for FF evaluation if the mouse remained in the same state across atleast 10 trials for the complete duration of the time-window (150 ms). For all analyses, FF was evaluated only on units whose receptive fields were at least 20 degrees away from the monitor’s edge (Figures 4**F**, 6**F**).

### Mutual information

Mutual information (MI) measures the reduction in uncertainty about one random variable when the value of another variable is known (Cover 1991). For two variables, *X* and *Y*, it is calculated as the difference between the total entropy of *X*, denoted as *H*(*X*), and the entropy that remains in *X* after learning the value of *Y*, referred to as the conditional entropy *H*(*X|Y*).

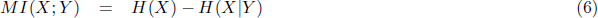

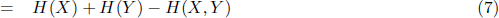

Similar to correlation, MI is symmetric in *X* and *Y*, meaning that *MI*(*X*; *Y*) = *MI*(*Y* ; *X*). This is evident when MI is re-written in terms of joint entropy between the variables (equation 7). However, MI surpasses correlation in its capacity to capture non-linear connections between variables. Given that responses of visual neurons can be highly non-linear functions of the visual input, we favored MI as our primary metric to quantify the amount of pixel-level information embedded in the neuronal activity of the visual cortex. Yet, calculating entropy requires knowledge of the joint probability distribution function (pdf) of the random variables, which is often unavailable. Many studies resort to ‘plug-in’ estimators that involve intricate evaluations of individual pdfs, a particularly onerous task for sizable datasets like ours. To sidestep the need for pdf estimation, we employed a matrix-based entropy estimator whose properties have been shown to align with the axiomatic properties of Renyi’s *α*-order entropy (*α >* 0) (Sanchez Giraldo et al. 2015).

Here, we provide a brief description of the process of entropy evaluation using the estimator, for specific details see Sanchez Giraldo et al. 2015. First, the sample variable, *X* = [*x*1*, x*2*, …, xN*] *∈* R*^N×M^*, is projected into a reproducing kernel Hilbert space (RKHS) through a positive definite kernel, *κ* : *X × X 1→* R. Next, a corresponding normalized Gram matrix, denoted as *A*, is generated from the pairwise evaluations of the kernel, *κ*. In this matrix, each entry *A_ij_* is calculated as 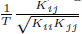, where *K* = *κ*(*x, x*) and *K ∈* R*^N×N^* . The entropy estimator then defines entropy using the eigenspectrum of the normalized Gram matrix *A*, following the equation (8), where *λ_j_* (*A*) represents the *j^th^* eigenvalue of matrix *A*. Finally, the joint entropy, *H*(*X, Y*) or *S_α_*(*A, B*), is evaluated as the entropy of the Hadamard product, *A ◦ B* (equation 9), where *B* is the normalized Gram matrix associated with *Y* . The Hadamard product is interpreted as computing a product kernel, *κ*((*x_i_, yi*), (*x_j_, yj*)).

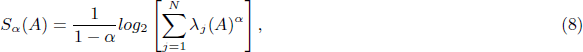

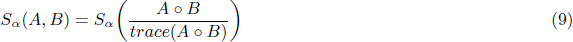

In our analyses, we consider *X* = [*x*_1_*, x*_2_*, …, x_N_*] *∈* R*^N×M^* to represent the spike count matrix for all neurons in the population, where each *x_i_ ∈* R*^M^* is a vector containing spike counts from *M* neurons at time *i*. Similarly, *Y* = [*y*_1_*, y*_2_*, …, y_N_*] *∈* R*^N×P^* is a matrix containing image pixels, with each *y_i_* representing a flattened vector of all pixels in the stimulus image at time *i*. For state-wise analyses of MI (Figure 4**G**), we exclusively considered times corresponding to the specific state under examination. MI was computed per trial, but only when the subject had spent at least 3 seconds in the particular state during the trial, i.e., *i ∈* [3, 30] seconds. Each frame was downsampled by a factor of 5, and spike counts were evaluated in 30 ms bins to match the stimulus frame rate. To constrain the metric between [0, 1], all MI measures were normalized by the geometric mean of the individual entropy of the two variables, *S_α_*(*A*) and *S_α_*(*B*) (Strehl and Ghosh 2002). The values presented in the paper are averages taken across all subjects (Figure 3**F**, 4**G**).

Entropy estimation is dependent on two hyperparameters: the order, *α*, and the kernel, *κ*. Given the sparsity of neural activity data, we chose the order, *α*, to be 1.01. Next, *κ* is a positive definite kernel that determines the RKHS and thus dictates the mapping of the probability density functions (pdfs) of the input variables to the RKHS. For our analyses, we employed a non-linear Schoenberg kernel (equation 10). These positive definite kernels are universal, in that, they have been proven to approximate arbitrary functions on spike trains (Park et al. 2013). The window, *w*, to evaluate spike counts was set to 30 ms to match the frame rate of the visual stimulus, and the kernel width, *σ_k_*, was determined using Scott’s rule (Scott and Sain 2005).

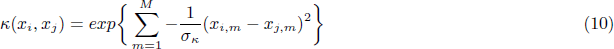

### Stimulus features

Capitalizing on the ethological significance of a naturalistic stimuli (Dan et al. 1996; Srinivasan et al. 1982; Yao et al. 2007) and to mitigate sudden changes in neural activity due to abrupt changes in visual stimulus, our analysis centered on neural data obtained from repeated viewings of a 30-second natural movie clip. We anticipated that the statistical properties of the clip would significantly contribute to explaining neuronal variability. In order to reveal any statistical preferences of neurons across the cortical hierarchy, we constructed stimulus features from both low- and high-order (*>* second-order moments) properties of the pixel distribution. The low-order features included image intensity and contrast, whereas, the high-order features included kurtosis, entropy, energy, and edges.

*Intensity and contrast:* These metrics captured the first and second order statistics of the image, and they were evaluated as the mean (*µm*) and standard deviation (*σ_m_*) of all the pixel values in each image frame, *I*, respectively.

*Kurtosis:* A higher-order statistic of the pixel distribution, Kurtosis measures the extent to which pixel values tend to cluster in the tails or peaks of the distribution. This metric was computed on the distribution of pixels within each image frame by determining the ratio between the fourth central moment and the square of the variance.

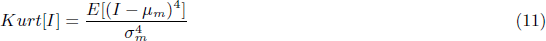

*Entropy:* To assess the average information content within each image frame, entropy was calculated based on the sample probabilities (*pi*) of pixel values spanning the range of 0 to 255.

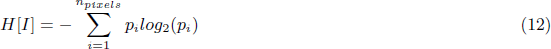

*Energy* : Similar to the quantification of face motion energy (Stringer et al. 2019), we evaluated image energy as the absolute sum of the differences between the pixel values of consecutive frames.

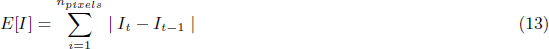

*Edges*: Given the observed line and edge selectivity of visual cortical neurons (Hubel and Wiesel 1968), we devised this metric to quantify the fraction of pixels that contribute to edges within a given image frame. For the identification of edges in each frame, we employed Canny edge detection (cv.Canny). This technique involves several sequential steps. First, a 2D Gaussian filter with dimensions of 5 *×* 5 pixels was applied to the image to reduce noise. Subsequently, the smoothed image underwent convolution with Sobel kernels in both horizontal and vertical directions, producing first derivatives along the respective axes, as described in equations (14 - 15). The resulting edge directions (Θ) were approximated to one of four angles: [0*^◦^,* 45*^◦^,* 90*^◦^,* 135*^◦^*]. To refine the edges, a process called edge thinning was used. During this step, the entire image was scanned to locate pixels that stood as local maxima within their gradient-oriented vicinity. These selected pixels moved on to the subsequent phase, while the rest were set to zero. Lastly, two threshold values were introduced for edge identification. Edges with intensity gradients below the lower threshold were disregarded, whereas those with gradients above the higher threshold were retained as ‘sure edges’. Pixels with gradient intensities falling between these two thresholds were analyzed based on their connection to a ‘sure’ edge. Ultimately, the output of the Canny edge detector was a binary image outlining the edge-associated pixels. The metric ‘edges’ was computed as the mean value of this binary image.

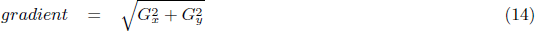

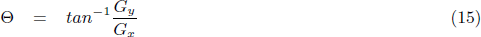

### Input data for the HMM-predictor model

Identical set of features were employed to predict both averaged neuronal population activity and single neuron responses. These features were grouped into three distinct categories to evaluate the respective contributions of each set of variables. The categorization of features is as follows: 1. stimulus features, 2. behavioral features, and 3. features encompassing internal brain dynamics, which included raw LFPs from the same cortical area, as well as averaged neuronal population activity from visual areas other than the target area. For raw LFPs, representative channels were once again selected across the cortical depth, ensuring the inclusion of one channel from each layer. Stimulus and behavioral features were sampled at a frequency of 30 Hz. However, to align with this temporal resolution, both LFPs and averaged population activity were binned into 30 ms bins, where each bin represented an average signal value within the respective time window.

The broad range of input features exhibited pronounced inter-correlations, and constructing an encoding model using a design matrix containing linearly dependent columns inherently jeopardizes model reliability. To avoid this multicollinearity in the design matrix, we systematically orthogonalized the input features using QR decomposition (Mumford et al. 2015). QR decomposition of a matrix, denoted as *M ∈* R*^m×n^*, yields *M* = *QR*, where *Q ∈* R*^m×n^* denotes an orthonormal matrix and *R ∈* R*^n×n^* represents an upper triangular matrix. Consequently, matrix *Q* spans the same space as the columns of *M*, ensuring that the columns of *Q*maintain mutual orthogonality. As QR decomposition systematically decorrelates each column from all preceding ones, the arrangement of columns within the matrix becomes pivotal.

Prior to constructing the time-shifted design matrix, we first orthogonalized internal brain activity relative to all other input features, positioning these columns towards the latter part of the matrix, *M* . This step was aimed at reducing the potential influence of stimulus and behavior features on brain activity (Musall et al. 2019). We retained the original definitions of stimulus features due to their limited correlations within and across groups (*r_within_* = 0.3 *±* 0.1, *r_across_* = 0.06 *±* 0.1, mean *±*std Figure 5**A**, panel 2). Given the strong correlations between behavioral features (*r_within_* = 0.4 *±* 0.2, *r_across_* = 0.07 *±* 0.07, mean *±* std), we applied QR decomposition to decorrelate all behavioral variables among themselves. The final collection of input features for the full model comprised behavioral features that had undergone orthogonalization among themselves, stimulus features in their original form, and internal brain activity features that were orthogonal both within and across the categories of features. Next, each input signal of length *τ* was organized such that each row consisted of variables shifted in time by one frame (30 Hz) relative to the original, also known as a Toeplitz matrix. Lastly, to structure the design matrix, the various input signals were time-aligned and concatenated. Including a time-shifted design matrix enabled us to account for the temporal dependency between various sources and neural activity. To determine the appropriate time dependency for each type of neural data (averaged neuronal population and single neuron activity), we tested a range of values (population model: [0.2 - 6]s, single-neuron model: [0.2 - 2]s) and chose the dependency that maximized the model’s cross-validated explained variation, *cvR*^2^ (Figure S7).

Lastly, when quantifying group-specific contributions using unique models, the features of internal brain activity were orthogonalized within the group. This approach was taken to prevent partial decorrelation across groups, as the designed stimulus features and behavioral features might not encompass the entire array of features encoded in neural activity. Such partial decorrelation could potentially obscure the interpretability of the contributions from each category of input features to spiking variability.

### HMM - predictor model

The linear HMM-predictor model was constructed to predict the averaged neuronal population activity and single-neuron spike rates. Unlike classical linear prediction models that assume constant relative contributions of various sources to spiking variability, the HMM-predictor model deviates from this assumption by accounting for variations in contributions resulting from internal state fluctuations. To achieve this, each predictor model learns regressors only from signals associated with a state. This approach enables us to delve into state-specific investigations of the relative contributions across the three distinct sources of variability outlined earlier. Each predictor model is tailored specifically to the neural activity in each state. Importantly, it should be highlighted that the HMM states are held constant. In other words, the HMM model is not optimized to improve predictions but maintains its established definitions based on LFPs. To quantify the contributions of the variability sources to the averaged population activity, we used ridge regression, whereas spiking activity was modeled using a generalized linear model (GLM).

#### Population model

To mitigate overfitting, the population model was trained with ridge regression. Ridge regression extends the cost function of ordinary least squares by introducing an additional *l*_2_ penalty, (*λ*), on the regression coefficients (*β*). This penalty effectively shrinks the coefficients of input variables that contribute less to the prediction, promoting smoother and more generalizable regression coefficients (equation 16). In our HMM based regression model, the design matrix *X_s_* and the regressand, *y_s_*, are informed by the HMM, comprising signals corresponding to one of three identified states (*s* = [*S_H_, S_I_, S_L_*]). The magnitude of the regularization penalty, *λ*, for weights in each state were individually determined through three-fold cross-validation of *R*^2^ on a randomly selected 30% subset of the dataset.

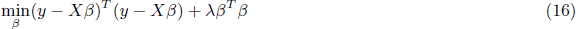

#### Single neuron model

A regularized Poisson GLM was used to model the firing rate of each neuron while taking into account variances associated with internal state fluctuations. The encoding model describes spike counts of single neurons as a Poisson distribution whose expected value can be modeled as the exponential of the linear combination of input features, i.e, *E*(*y|X*) = *e^θT X^* . The coefficients of the regression model, *θ*, are then estimated by penalized maximum likelihood with an *l*_2_ penalty on the coefficients (equation 17) (Pillow et al. 2008). To avoid overfitting, the magnitude of the regularization penalty, *λ*, for weights for each neuron in each state were individually determined using nested-five-fold cross-validation of *R*^2^ during training (Cawley and Talbot 2010).

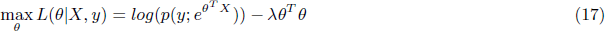

The final evaluation of the reported scores (Figures 5, 6) includes a five-fold cross-validation of explained variance (*cvR*^2^, equation 18), where *ŷ* is the predicted spike rate and *ȳ* is the mean of the true spike rate. The *cvR*^2^ values in Figure 6 were computed on spike counts of single neurons smoothed with a 50-ms Gaussian for each trial.

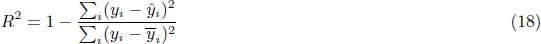

To quantify the state-wise contributions of the input features, we partition the dataset into training and testing sets such that each fold contains an equitable representation of signals from every state. This step was crucial to prevent any potential biases in estimating contributions due to an imbalance in the number of data points in each state. State-specific contributions were evaluated on the respective performance of the state-wise regressors, while overall performance was evaluated by concatenating the predictions across the three state models in each fold.

## Acknowledgements

Primary funding for this project was provided by the Allen Institute phase IV grant for Mindscope and Tsinghua-Peking Center for Life Sciences (C.L.S.). We thank the National Natural Science Foundation of China (92370116 to X.J.), and Tsinghua University Initiative Scientific Research Program for additional funding. We thank Ruben, Coen-Cagli, Ramakrishnan Iyer, Scott Linderman, Lukasz Kusmierz, Eric Shea-Brown, Nick Steinmentz and Ying Zhou for helpful discussions; Linzy Casal for assistance with planning and budgeting of the project; Benjie Miao for technical support.

## Author contributions

Conceptualization: X.J., S.A. Supervision and funding acquisition: X.J. Investigation and formal analyses: S.A., X.J. Validation and methodology: S.A., X.J., P.L., J.H.S., S.R.O., M.A.B, D.D. Software and visualization: S.A., X.J., S.R.O. Data collection: S.D., H.B., X.J., J.H.S. Original draft written by S.A., X.J., with input and editing from S.R.O., J.H.S, P.L., M.A.B, D.D., C.K., H.B., S.D. All co-authors reviewed the manuscript.

## Supplementary Figures

**Figure S1:**
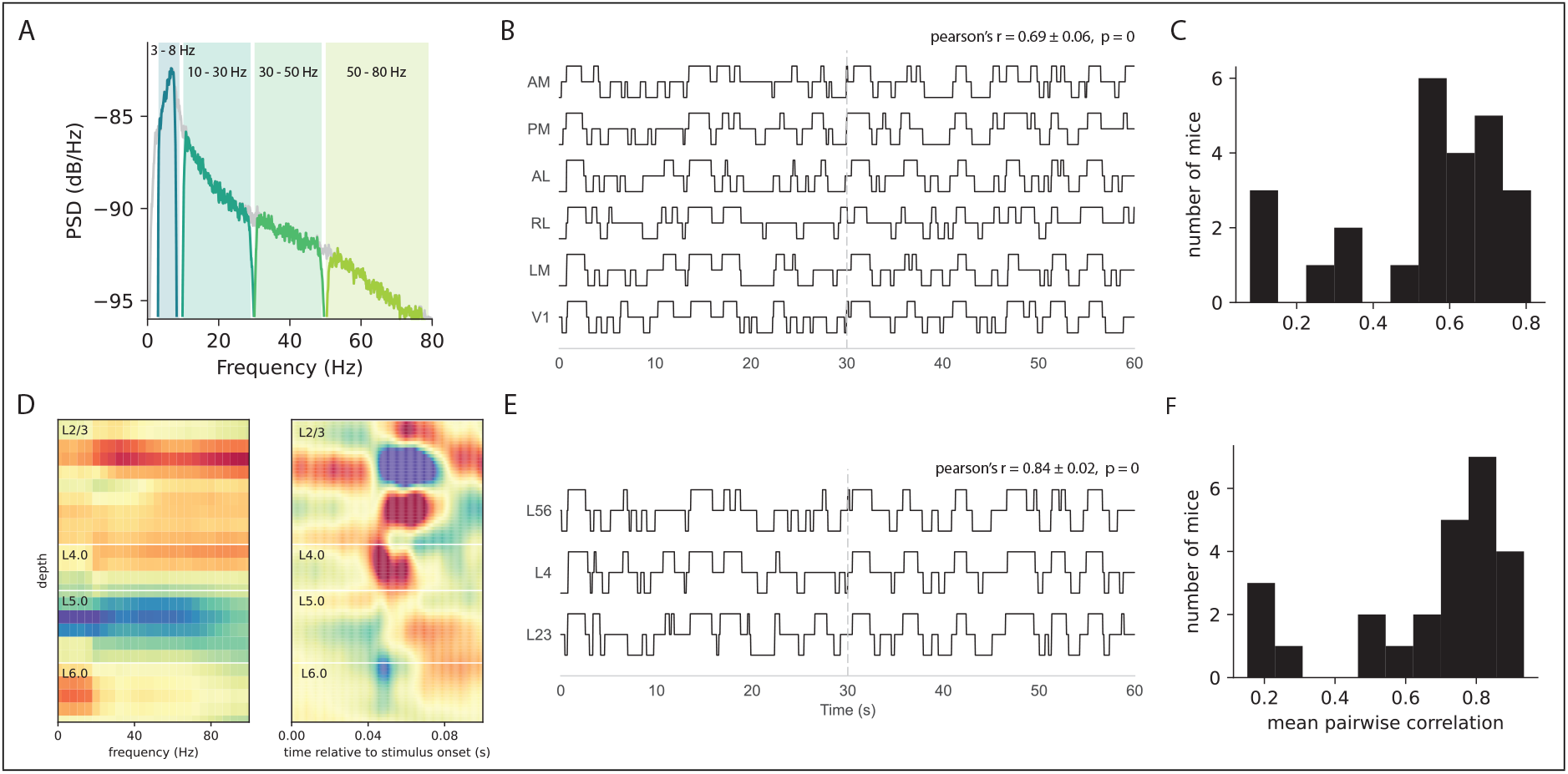
**A**, Complete power spectral density (PSD) in a single channel overlaid with the PSDs of the filtered LFPs in the respective frequency bands. **B**, State sequences estimated using Local field potentials (LFPs) from individual areas of an example mouse. One channel from each layer was incorporated into the Hidden Markov Model (HMM) input matrix. **C**, Histogram summarizing the average pairwise correlations between state sequences estimated from individual areas. **D**, Channel classification into L2/3, L4, or L5/6 based on analyses of the power spectral density (left) and current source density (right) of the LFPs along the cortical depth during the presentation of flashes. **E**, State sequences estimated using LFPs from individual layers of an example mouse. LFPs from all areas were included in the HMM input matrix. **F**, Histogram illustrating the average pairwise correlations between state sequences estimated from individual layers.

**Figure S2:**
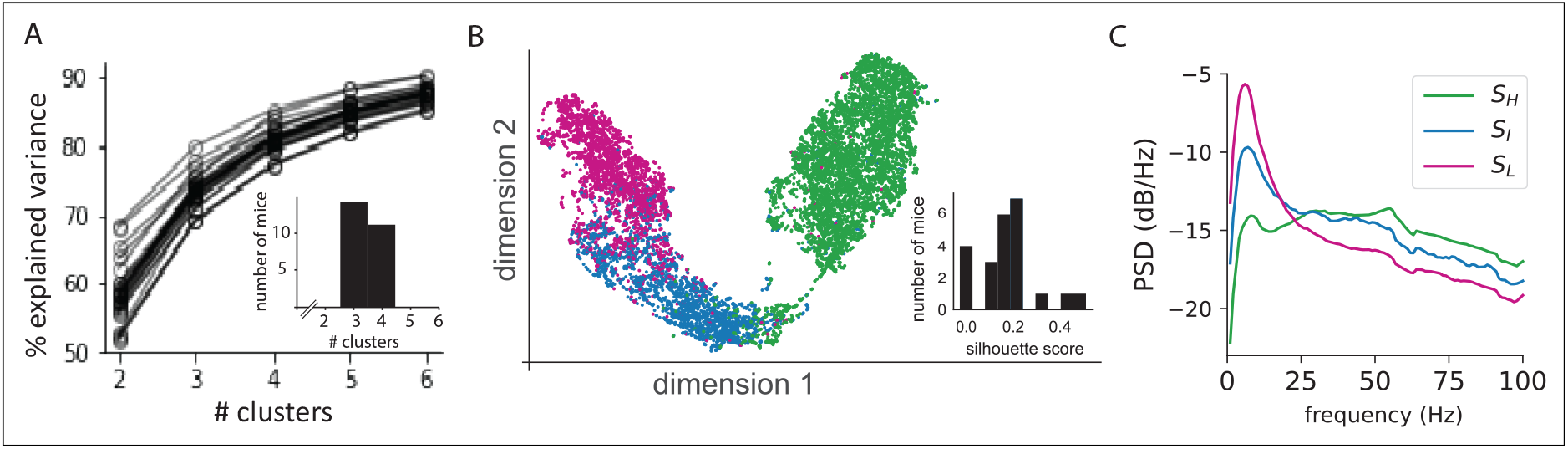
**A**, K-means clustering to validate the optimal number of states for the Hidden Markov Model (HMM). Elbow method on the variance explained by K clusters. (Inset) Histogram of the optimal number of states across all mice **B**, UMAP projection of the LFP inputs provided to the HMM in an example mouse. (Inset) Silhouette scores based on HMM states and UMAP projection. **C**, State-specific power-spectral density of all LFPs in V1 in an example mouse. Such decomposition in all mice further confirmed the spectral distinction observed across the different oscillation states.

**Figure S3:**
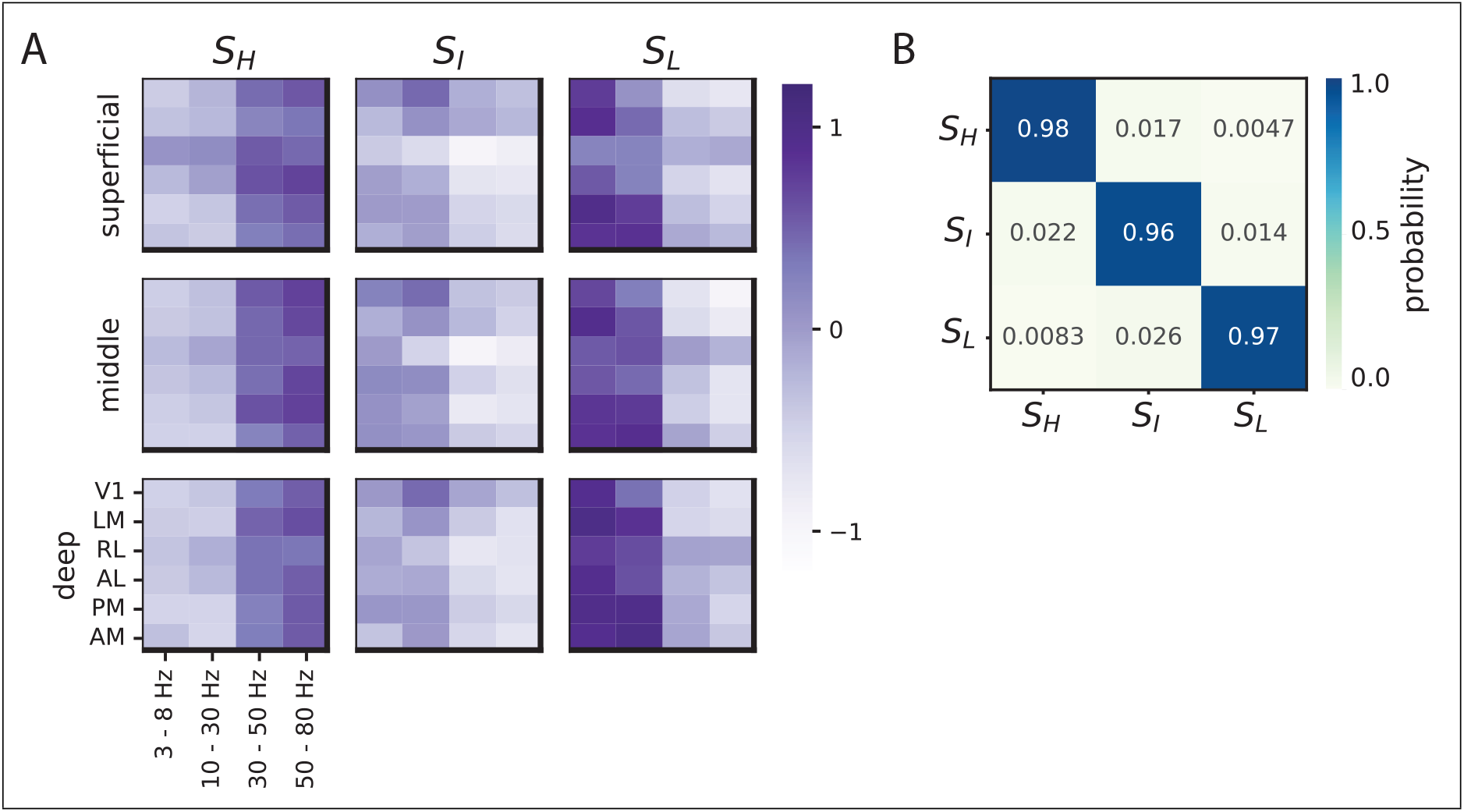
**A**, State emission matrix summarizing the means of each input features within the HMM. **B**, State transition probability matrix. Results from an example mouse.

**Figure S4:**
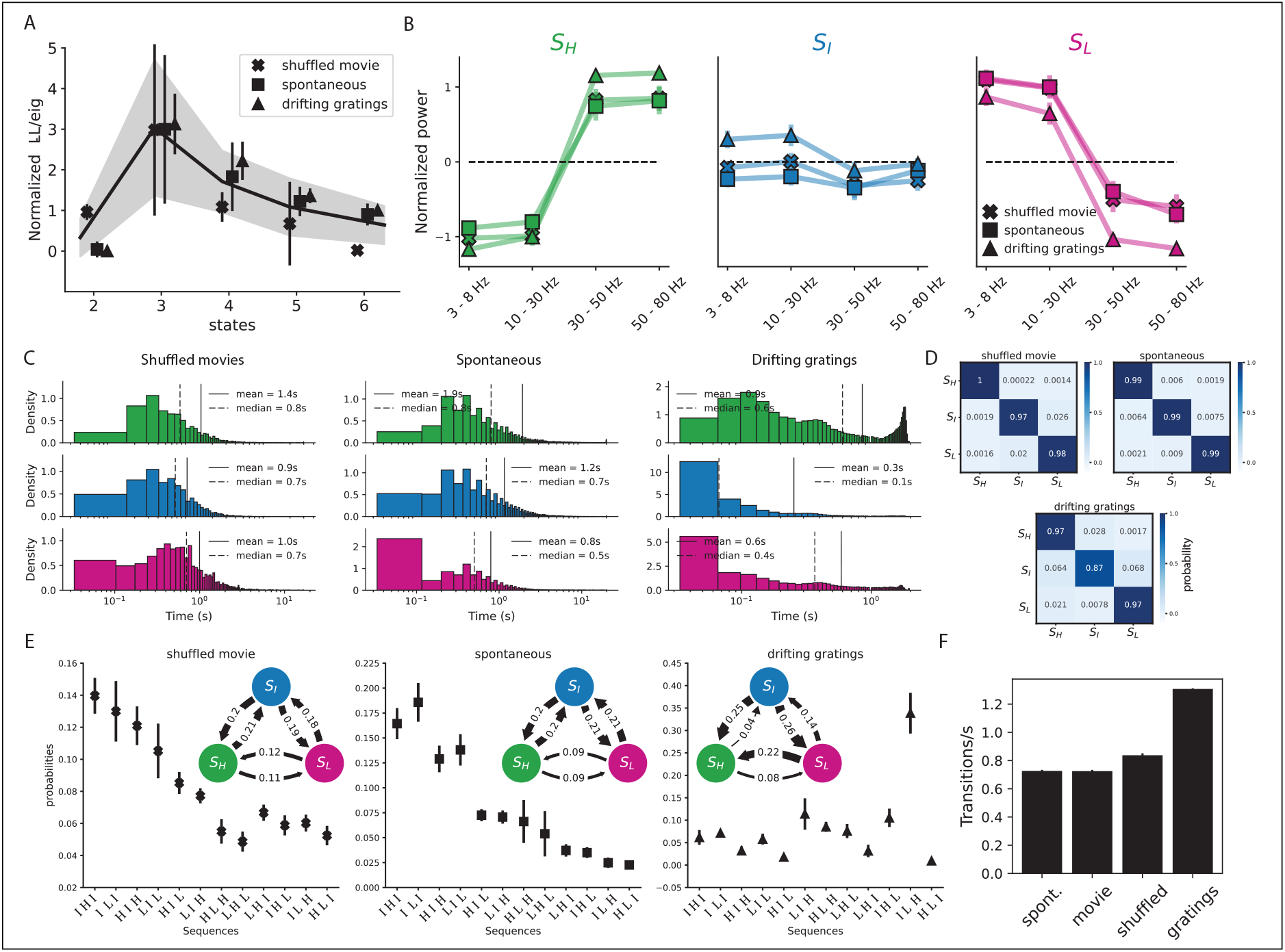
**A**, Model comparison among Hidden Markov Models (HMMs) across a range of latent states for different stimulus types. **B**, Distribution of LFP power in the three-state model as subjects viewed different stimuli. **C**, Dwell times in each state as subjects viewed various stimuli. **D**, Matrices depicting state transition probabilities. **E**, Average probability of observing 3-step or 2-step (inset) transition sequences to different states while viewing various stimuli. Transition probabilities were calculated from observed sequences averaged across all mice. **F**, Number of state transitions per second during the viewing of different stimuli.

**Figure S5:**
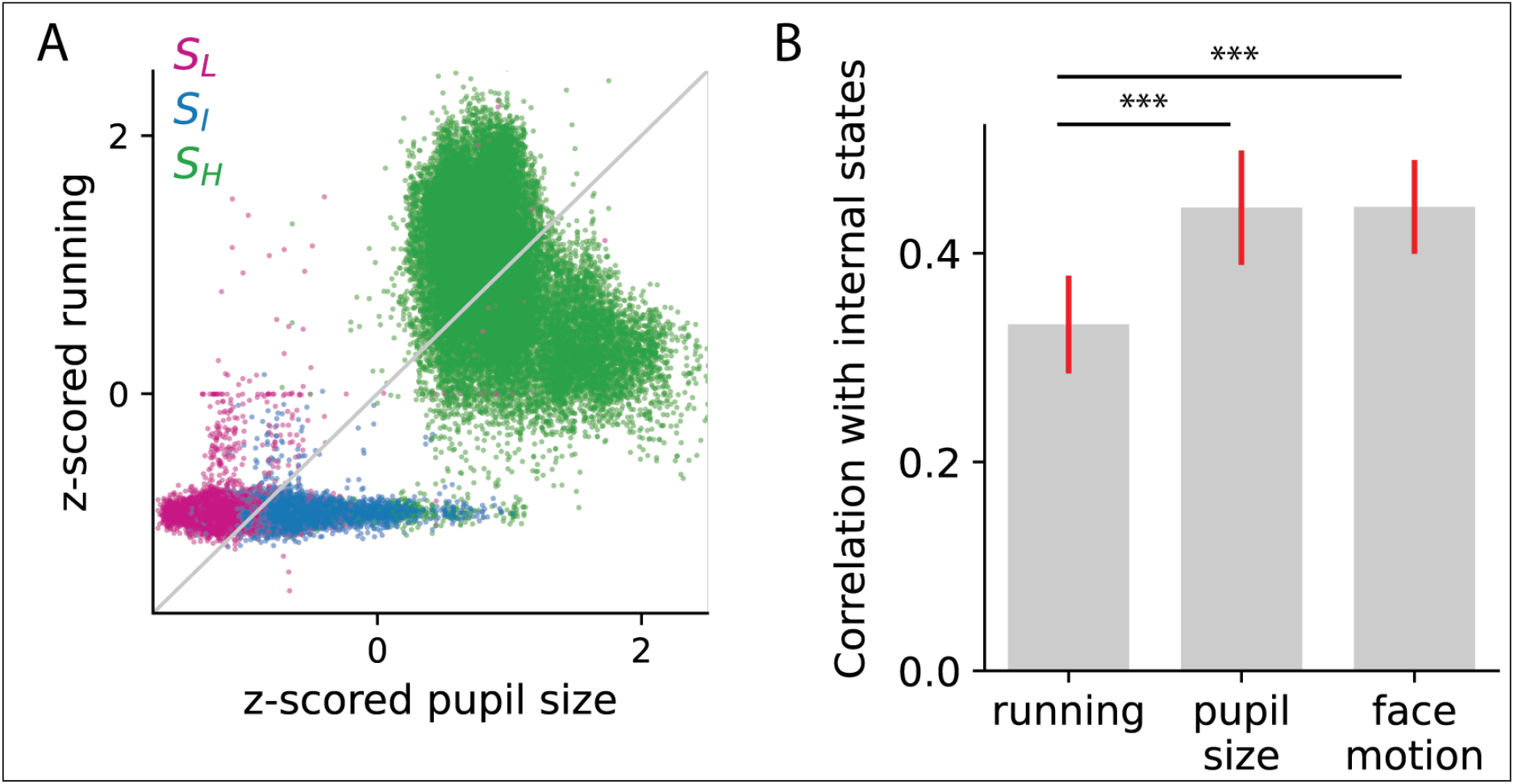
**A**, Scatter plot of pupil size and running speed color-coded to demarcate the time points of different states. **B**, Average correlation between behavioral states identified individually using running speed, pupil size and facial motion with internal oscillation states.

**Figure S6:**
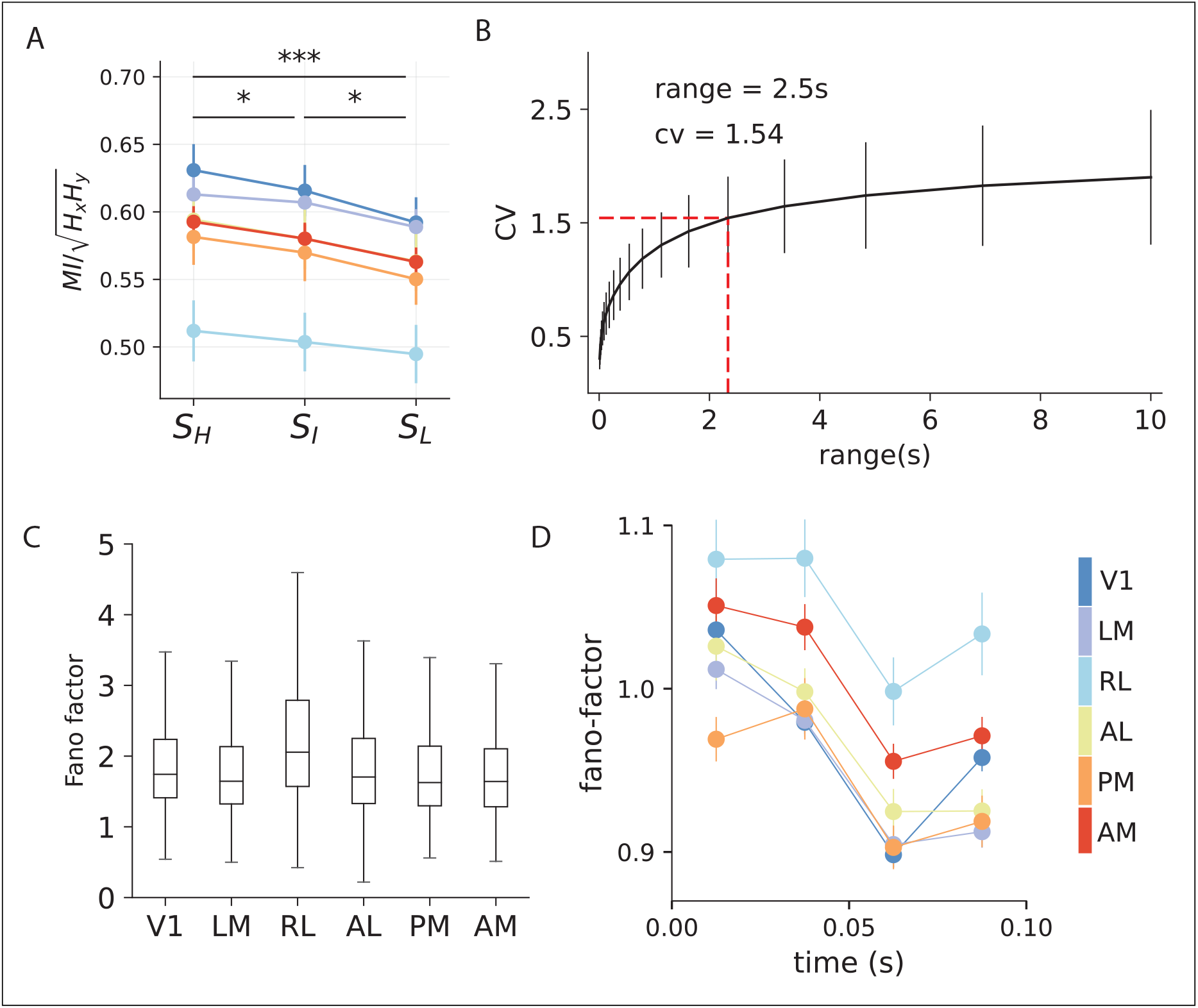
**A**, Information encoding along the visual hierarchy across all oscillation states, quantified using mutual information (MI). Error bars represent s.e.m. **B**, Time-scale estimation for the construction of inter-spike-interval histograms, utilized in the estimation of the coefficient of variation metric. **C**, Box plot summarizing Fano factors in each area (Pearson correlation with anatomical hierarchy scores excluding RL, *r_p−RL_* = *−*0.7*, p_p−RL_* = 0.11) **D**, Comparison of Fano factor across visual areas evaluated over time when the mice were exposed to full-field light flashes.

**Figure S7:**
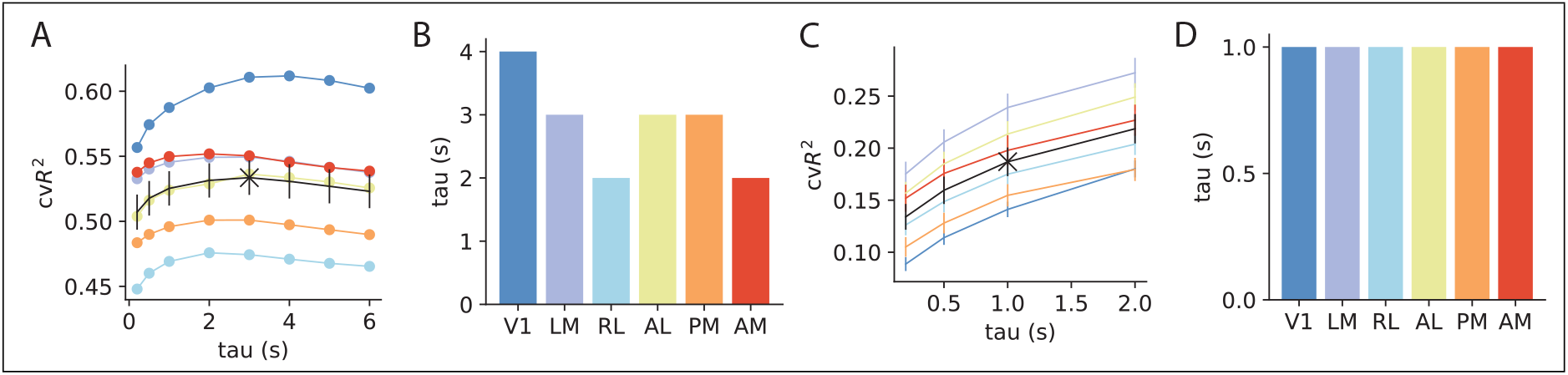
**A**, Selection of kernel length, *τ*, for HMM-regression model to predict variance in the averaged neuronal population activity. The kernel length, which had the maximum predictive power, was chosen. **B**,Optimal kernel length for area-wise HMM-regression models. **C**, Selection of kernel length, *τ*, for HMM-GLM model to predict single neuron variability. Kernel length was selected on cross validated *r*^2^ using the elbow method. Results from an example mouse. **D**, Optimal kernel length for area-wise HMM-GLM models for the example mouse.

**Figure S8:**
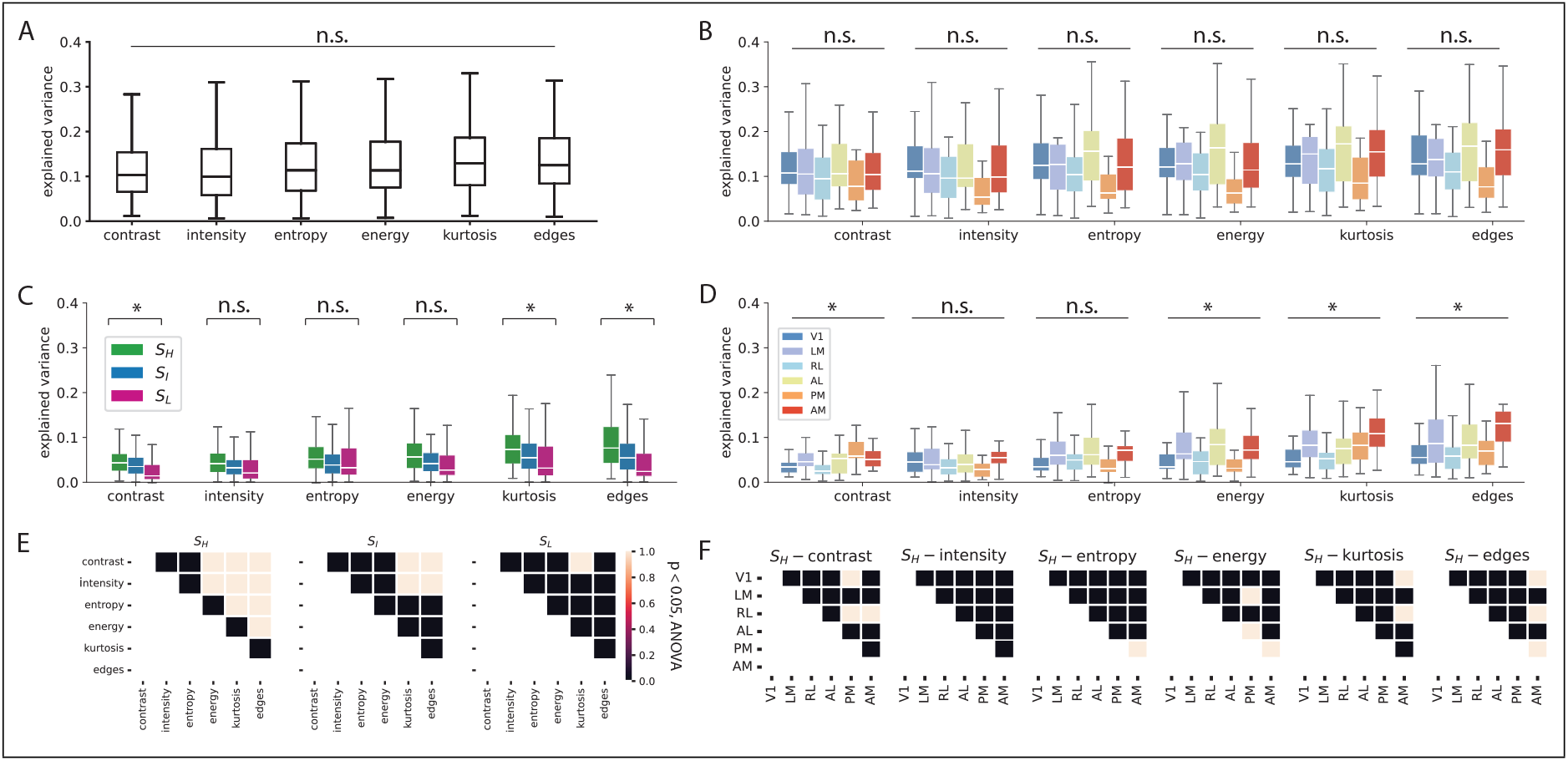
**A**, Summary of the variance explained in averaged population activity by different stimulus features. **B**, The contribution of different stimulus features to the variance of averaged population activity across visual areas. **C**, State-wise contributions of different stimulus features to averaged population activity. **D**, Same as **B**, but during the high-frequency state.**E**, Significance results for **C**, p *<* 0.05, corrected for multiple comparisons. **F**, Significance results for **D**, p *<* 0.05, corrected for multiple comparisons.

**Figure S9:**
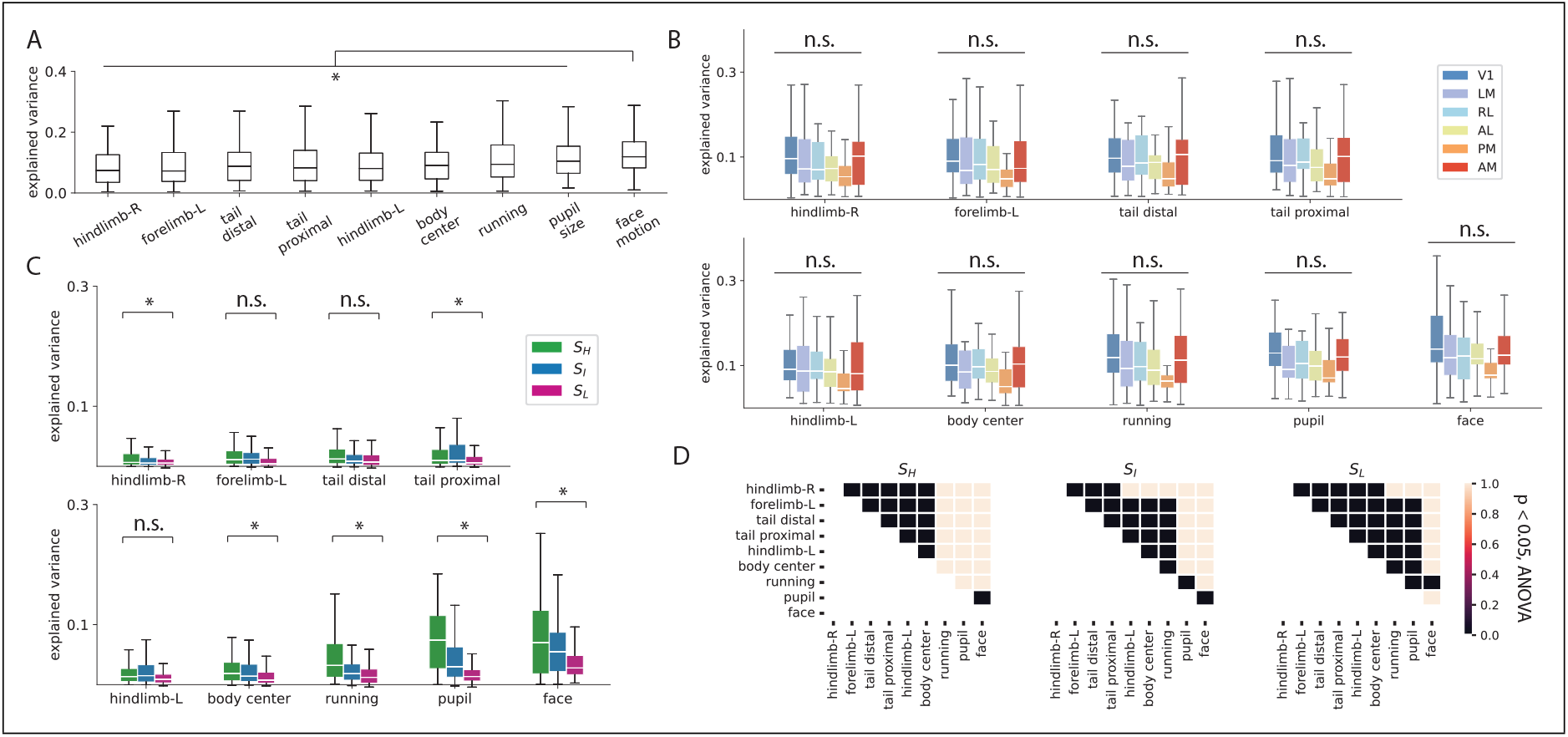
**A**, Summary of the variance explained in averaged population activity by various behavioral features. **B**, The contribution of different behavioral features to the variance of averaged population activity across visual areas. **C**, State-wise contributions of behavioral features to averaged population activity. **D**, Significance results for **C**, p *<* 0.05, corrected for multiple comparisons.

**Figure S10:**
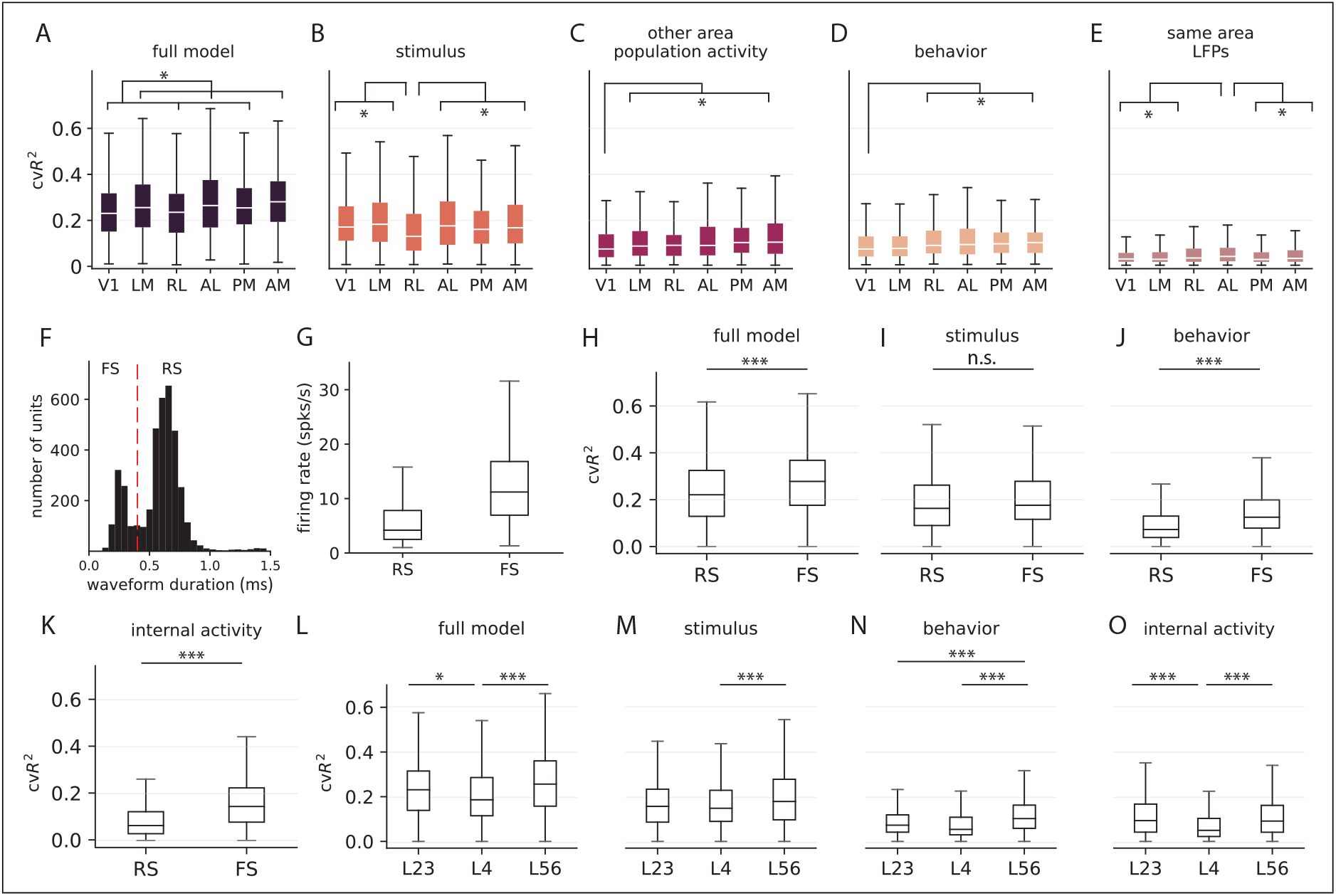
**A-E**, Contributions from different input features to explain single neuron variance in different visual areas. Analysis of the relative contributions to area-specific single-neuron variability showed that the anterolateral visual areas (LM, AM, and AL) had the highest explained variance of approximately 26.2*±*0.9% (mean *±* std). Consistent with other results, neurons in RL did not encode stimulus features as well as the other visual areas. However, behavior and LFP features explained the most variance in RL and AL neurons 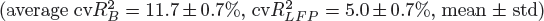, while these features were the least predictive of activity in V1 neurons 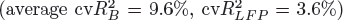. The predictive power of the averaged neuronal population activity from neighboring areas had trends similar to that observed in the population model, with V1 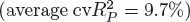 neurons being the least predictive and AM the most predictive 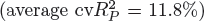. Moving across layers, L4 neurons reported the least explained variance, while deep-layer neurons consistently had the highest explained variance across all categories of input features (Figure S10**L - O**). **F**, Classification of single units into regular spiking (RS) and fast spiking (FS) based on waveform duration. **G-K**, Contributions from different input features to explain single neuron variance across RS and FS cell-types. **L-O**, Contributions from different input features to explain single neuron variance across the cortical depth.

